# Adaptive mechanisms of social and asocial learning in immersive collective foraging

**DOI:** 10.1101/2023.06.28.546887

**Authors:** Charley M. Wu, Dominik Deffner, Benjamin Kahl, Björn Meder, Mark H. Ho, Ralf H.J.M. Kurvers

**Affiliations:** Human and Machine Cognition Lab, University of Tübingen, Tübingen, DE; Centre for Adaptive Rationality, Max Planck Institute for Human Development, Berlin, DE; Department of Computational Neuroscience, Max Planck Institute for Biological Cybernetics, Tübingen, DE; Excellence Cluster: Science of Intelligence, Technical University Berlin, DE; Department of Psychology, University of Marburg, Marburg, DE; Institute for Mind, Brain and Behavior, Department of Psychology, Health and Medical University, Potsdam, DE; Department of Psychology, New York University, New York, NY

## Abstract

Human cognition is distinguished by our ability to adapt to different environments and circumstances. Yet the mechanisms driving adaptive behavior have predominantly been studied in separate asocial and social contexts, with an integrated framework remaining elusive. Here, we use a collective foraging task in a virtual Minecraft environment to integrate these two fields, by leveraging automated transcriptions of visual field data combined with high-resolution spatial trajectories. Our behavioral analyses capture both the structure and temporal dynamics of social interactions, which are then directly tested using computational models sequentially predicting each foraging decision. These results reveal that adaptation mechanisms of both asocial foraging and selective social learning are driven by individual foraging success (rather than social factors). Furthermore, it is the degree of adaptivity—of both asocial and social learning—that best predicts individual performance. These findings not only integrate theories across asocial and social domains, but also provide key insights into the adaptability of human decision-making in complex and dynamic social landscapes.

## Introduction

Humans have a unique capacity for social learning that differentiates us from other animals^1,2^. We are remarkably flexible in how we learn from others^3–5^, dynamically integrate personal and social information^6–8^, and selectively favor social learning when our own capabilities seem lacking^9,10^. And while a number of recent studies have begun to bridge individual and social decision-making^7,11–13^, they either assume fixed strategies or arbitrary mixtures of social and asocial learning. Thus, we still know very little about the mechanisms driving adaptation to different environments and circumstances, allowing us to dynamically arbitrate and integrate both asocial and social learning strategies^14–16^.

Historically, research on asocial and social learning has progressed largely independently from one another. Theories of asocial learning typically assume that decision makers operate alone in a vacuum^17,18^, while theories of social learning^19–21^ often greatly simplify—or entirely omit—individual learning mechanisms. Early work investigated the trade-off between individual and social learning through the lens of the producer vs. scrounger dilemma^22–25^, assuming either pure individual learning (i.e., producing) or pure social learning (i.e., scrounging)^26^. In this setting, scrounging comes at the cost of reduced opportunities for producing, with any strategy having *frequency-dependent fitness*, meaning one’s performance depends on the ratio of strategies in one’s group. This dynamic is illustrated in Roger’s Paradox^27^, where too many imitators in a group leads to both lower individual and group fitness. While theoretical models often show that an intermediate balance of social and asocial learners leads to the best outcomes^28,29^, it is still largely unknown how people dynamically negotiate this balance under realistic conditions and how they adapt to different environmental contexts. For instance, whether adaptation is driven by asocial or social cues, and whether these mechanisms operate independently or interactively with one another. Modeling dynamic strategy selection in social contexts is particularly difficult, because the availability and quality of social information constantly changes as result of both individual decisions and group dynamics^30^. Thus, this gap represents both theoretical and empirical challenges, requiring new methods to capture the complex and dynamic nature of human adaptability, which we seek to address in this current study.

Here, we use a collective foraging task programmed in an immersive Minecraft environment (Fig. 1a-d) to study how people adapt their asocial and social learning strategies to different resource distributions (random vs. smooth; Fig. 1e) and to different dynamic contexts (e.g., individual performance and social observations of success). The virtual environment imposes a limited field of view, creating a natural trade-off between allocating visual attention towards individual search or towards peers for social learning (in contrast to REFs^31,32^). Using a novel method for automating the transcription of visual field data (Fig. 1c; see Methods), we can identify which participants and which elements of the environment were visible at any point in time. This allows us to dynamically integrate visual attention with spatial trajectories and foraging decisions, providing a common framework for studying the drivers of adaptive behavior.

**Figure 1.**
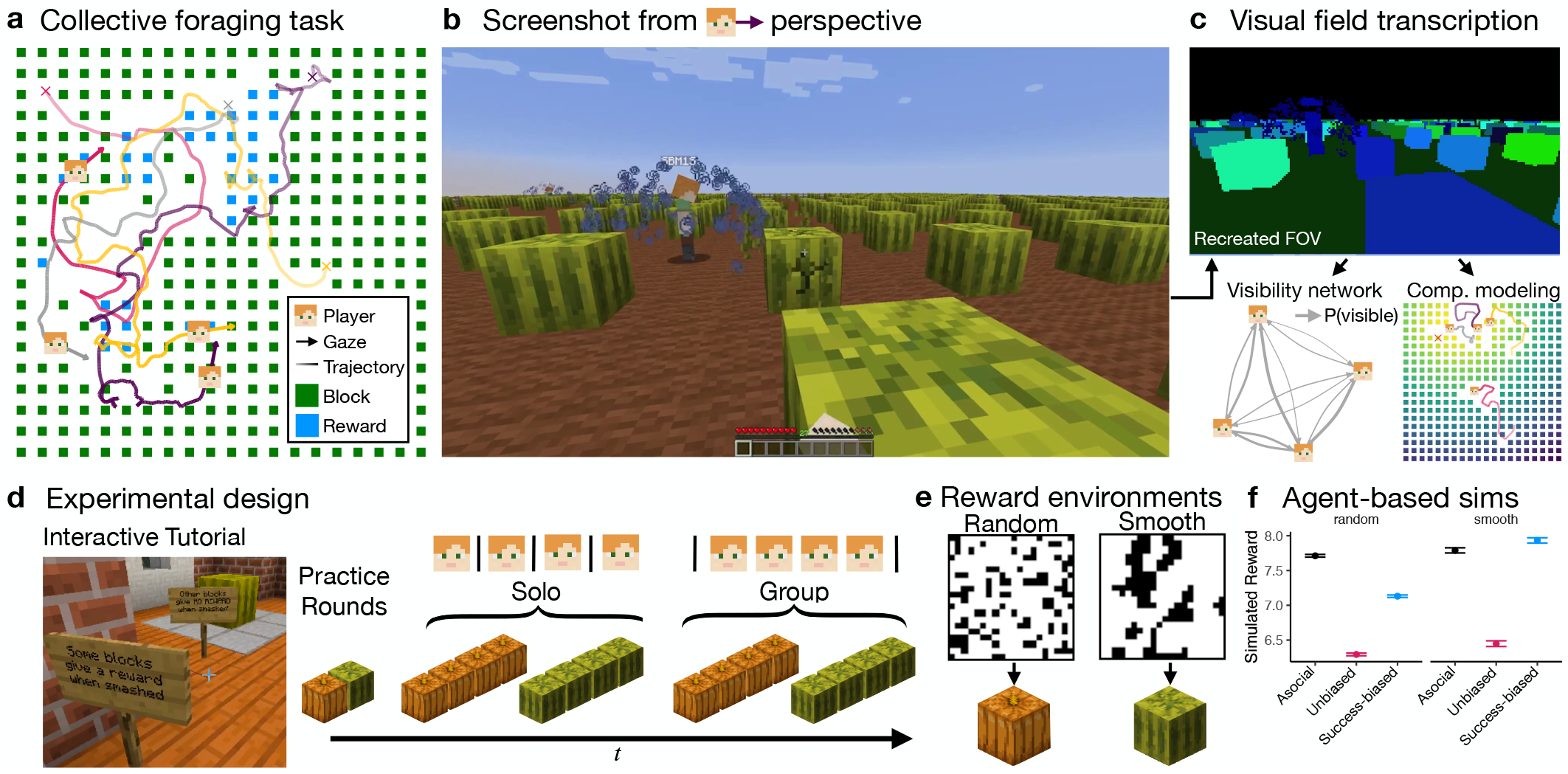
Collective foraging task implemented in the Minecraft game engine. (**a**) Participants foraged for hidden rewards in a field with 20×20 resource blocks. Each round took 120 seconds, with players starting from random locations (crosses) and heading directions (arrows). (**b**) Screenshot from a player’s perspective. Rewards (blue splash) are visible to other players, providing relevant social information for predicting nearby rewards in smooth—but not random—environments (Panel e). (**c**) Automated transcription of each player’s field of view (FOV) used in visibility and model-based analyses (see Methods). Participants learned about the task in an interactive tutorial (Supplementary Video 1) before completing two practice rounds. The main experiment consisted of 16 rounds (counterbalanced order), manipulated across condition (solo vs. group) and reward structure (random vs. smooth) with four consecutive rounds of the same type (Supplementary Videos 2-4). (**e**) Random environments had uniformly sampled rewards, while smooth environments had spatially clustered rewards. Each black pixel indicates a reward from a representative sample, with both environments having the same base rate *p* (reward) .25. The mapping to pumpkins and watermelons were counterbalanced between sessions. (**f**) Agent-based simulations (see Methods) show a benefit for success-biased social learning over asocial learning in smooth, but not random environments, whereas unbiased social learning performs poorly in both. Dots show the mean and error bars show the 95% CI over 10k simulations. This study is not approved by or associated with Mojang or Microsoft. Screenshots and images are used according to the Minecraft usage guidelines.

Adaptive mechanisms have been independently studied in both asocial foraging and social learning, however the two approaches have yet to be integrated in a single framework^15^. In asocial foraging, *area-restricted search*^33^ (ARS) has been used to describe an adaptive search strategy from species as diverse as bacteria^34^ to humans^35^, where the locality of search is modulated by foraging success: rich rewards drive local search, while poor rewards promote increased search distances. Although ARS is able to account for highly adaptive search patterns, it has yet to be integrated with social learning^15^. Adaptive mechanisms have also been proposed in social settings, based on context-dependent strategies that compare the quality of individual vs. social information^4,9,10^. Enquist et al.^4^ proposed two adaptive strategies: a critical social learner that first tries social learning, but switches to individual learning if social learning proves unsatisfactory, and a conditional social learner, that conversely tries individual learning first, but switches to social learning if necessary. While more flexible than strategies with a fixed level of social learning^27^, these approaches still lack an account of the selectivity of social learning with respect to whom to learn from^5,36^ and have yet to be integrated with asocial foraging^15^ and reward prediction mechanisms^37^. Here, we bridge this gap through integrative behavioral and model-based analyses.

In this study, we combine visual field analysis with highresolution spatial trajectories to provide an integrative perspective on how asocial and social learning mechanisms complement one another in a dynamic and integrative fashion. Our results show that people dynamically *adapt* asocial and social learning mechanisms to both the environment and individual performance and *selectively* direct social learning towards successful individuals (Fig. 2). Our behavioral analyses capture both the structure and temporal dynamics of social interactions (Fig. 2-3), which are then directly tested using computational models sequentially predicting each foraging decision (Fig. 4). Our winning model integrates adaptive mechanisms of asocial and social learning under a single framework, revealing that individual success (rather than social factors) drives changes in asocial foraging patterns and increases both the amount of social learning and selectivity towards successful individuals. Furthermore, it is the degree of adaptivity—of both asocial and social learning—that best predicts individual performance.

**Figure 2.**
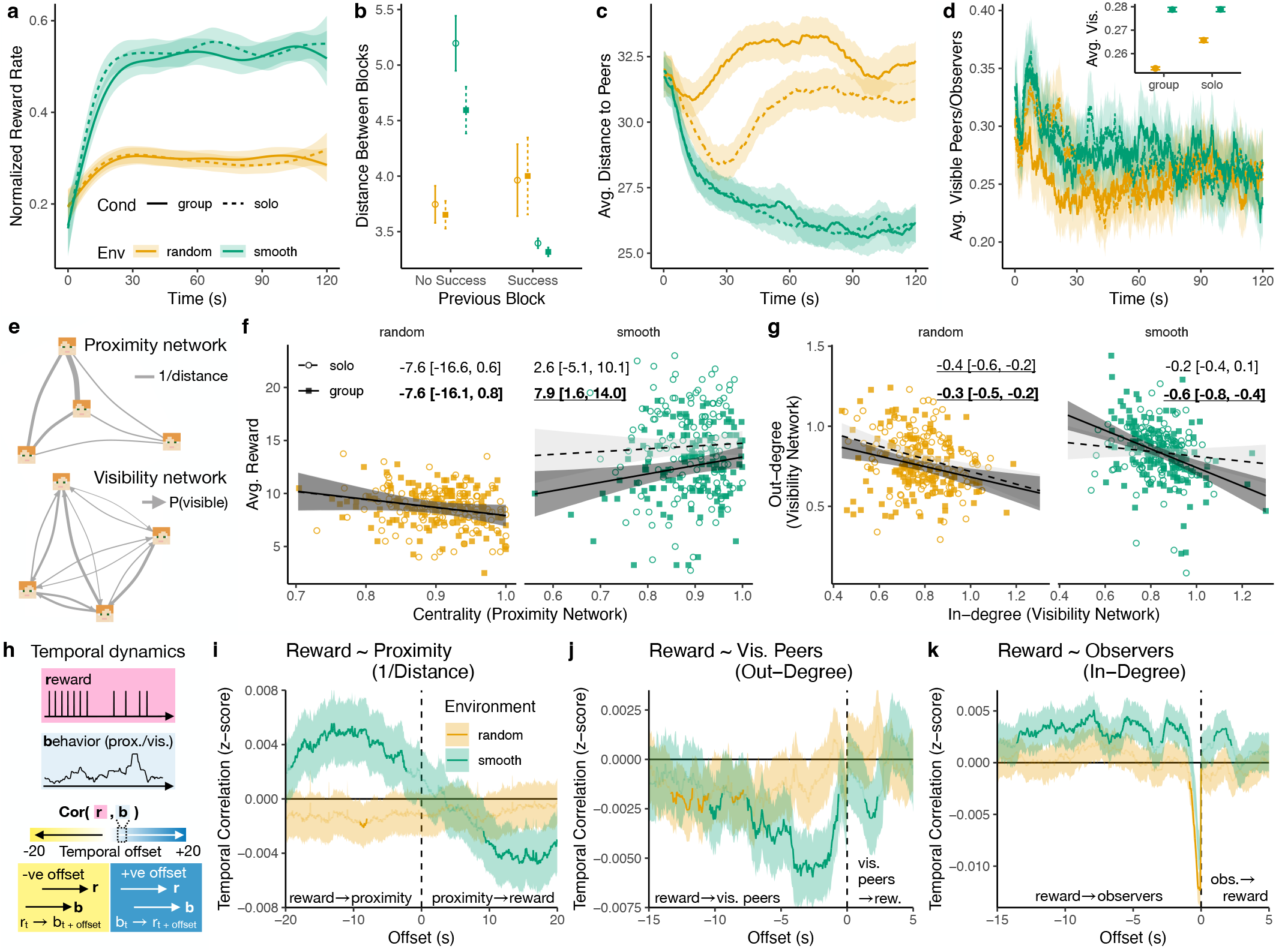
Behavioral results. **a**) Normalized reward rate controlling for reward depletion (see Fig. S1). Lines are from a generalized additive model (GAM) for the smoothing of binary data (group means and 95% CI). All CIs are over the *n =* 128 participants. **b**) Foraging distance as a function of previous success. Dots show the group means, while error bars indicate the 95% CI. **c**) Average distance to peers over time (see Fig. S5a for regression results), with lines showing aggregate means (ribbons are 95% CI). **d**) Average visibility, with the inset showing group means marginalized over time (see Fig. S5b for regression). Both ribbons and error bars indicate 95% CI. Note that at the aggregate level, avg. visible peers (outbound) and avg. observers (inbound) are equivalent. (**e-g**) Network analyses. Panel **e** shows examples of proximity and visibility networks. Panel **f** shows average reward as a function of Eigenvector centrality computed on the proximity network. Each dot represents one participant; lines and ribbons show the fixed effect of a hierarchical Bayesian regression (± 95% HPDI), with text labels reporting the effect (group rounds in bold), where reliable effects (not overlapping with zero) are underlined. Panel **g** shows the correspondence between in- and out-degree of the visibility network. Each dot represents one participant; the regression line is the fixed effect of a hierarchical Bayesian regression (Fig. S6). (**h-k**) Temporal dynamics of reward rate and proximity/visibility in group rounds (see Fig. S7 for comparison to solo rounds). Panel **h** provides a visual depiction of the analysis (see Methods for details). Results are shown in Panels **i**-**k**, where the y-axis shows the sign and strength of the correlation (after chance correction). Lines are group means and the ribbons show the 95% CI. Bold lines indicate significant clusters that survived a permutation analysis (see Methods). Effects at negative offsets indicate that rewards predicts future proximity/visibility, while effects at positive offsets indicate that proximity/visibility predicts future rewards. Positive effects indicate the rewards increased together with behavior, and vice versa for negative effects.

## Results

Participants (*n =* 128) foraged for hidden rewards either alone or in groups of four (solo vs. group; within-subject) in a virtual environment with 20 × 20 resource blocks (Fig. 1a-d). We manipulated the reward distribution (random vs. smooth) to modify the value of social learning (Fig. 1e). Smooth environments had clustered rewards, making social observations of successful individuals (visible as a blue splash; Fig. 1b) predictive of other rewards nearby. In contrast, random environments offered unpredictable rewards that provide no benefits for social learning.

Agent-based simulations (see Methods) support this intuition, with asocial learners dominating in random environments, whereas selective, success-biased social learners performed best in smooth environments (Fig. 1f). In contrast to non-competitive contexts^12,38^, where the social setting generally leads to better performance, here peers are both valuable sources of social information (in smooth environments) but also competitors for the same limited resources^3,39^ (Fig. S1). This emphasizes the need to adapt learning to a dynamically changing information environment, with similar real-world dynamics as developing marketplace innovations or engaging in scientific research^32,40,41^.

We start by exploring the influence of the environment and social information on behavioral patterns (Fig. 2a-d). However, only by analyzing the effects of network structure (Fig. 2e-g) and the temporal dynamics of social interactions (Fig. 2h-k) are we able to reveal the drivers of adaptive asocial and social learning. Next, we analyze social influence events (“pulls”) to ground our analyses in leader-follower dynamics (Fig. 3), which capture the active usage of social information through concrete changes in spatial position. Finally, we use computational models predicting sequential foraging decisions to directly test for different combinations of adaptive individual and social learning mechanisms (Fig. 4), integrating the rich spatial and visual dynamics of the task. All results are compared to an asocial baseline (using data from solo rounds), allowing us to specify which mechanisms are uniquely social phenomena.

**Figure 3.**
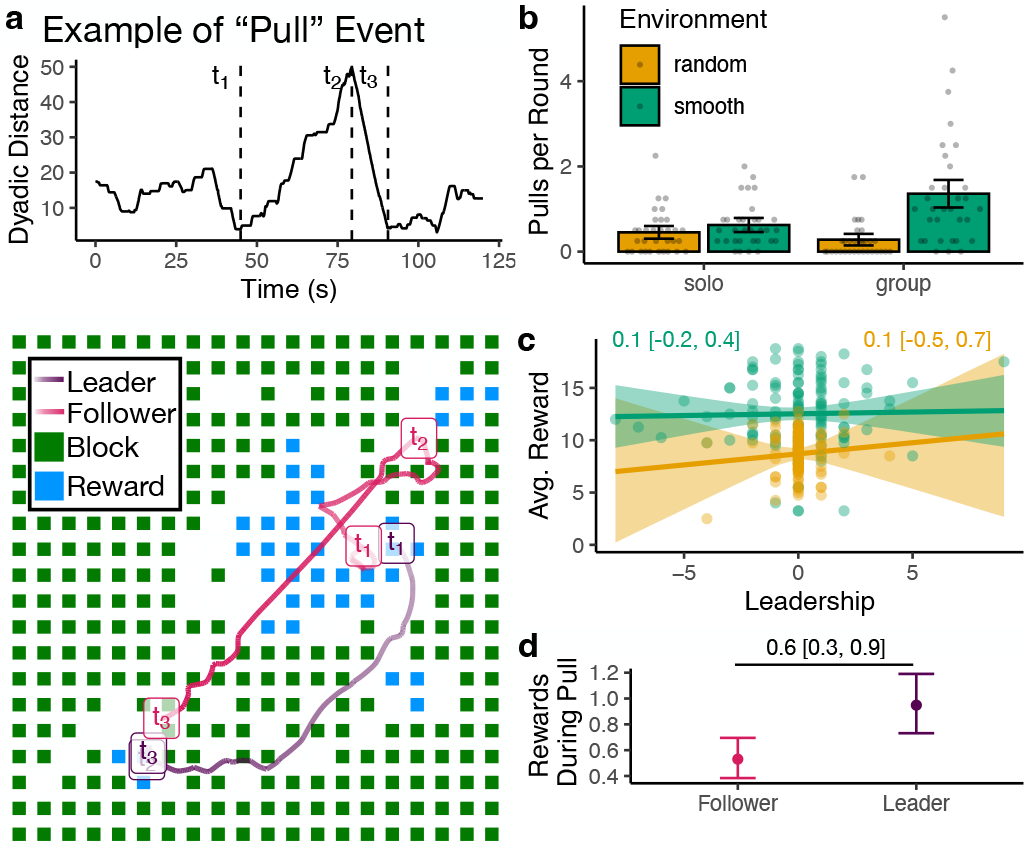
Social influence. (**a**) Example of a pull event, selected from min-max-min sequences in dyadic distance and filtered by a number of criteria (see Methods). The trajectories at the bottom are labeled with the three time points that define a pull, and show the state of the environment at time *t*_3_. Note that *t*_2_ for the leader largely overlaps with *t*_3_. (**b**) The average number of pull events per round (error bars indicate 95% CI and each dot is a session). We performed the same analysis on solo rounds as if participants were on the same field to provide an asocial baseline. (**c**) While leadership (*n*_leader_ − *n*_follower_) did not predict performance, (**d**) leaders had higher instantaneous rewards during pull events. Ribbons and error bars indicate 95% CI.

**Figure 4.**
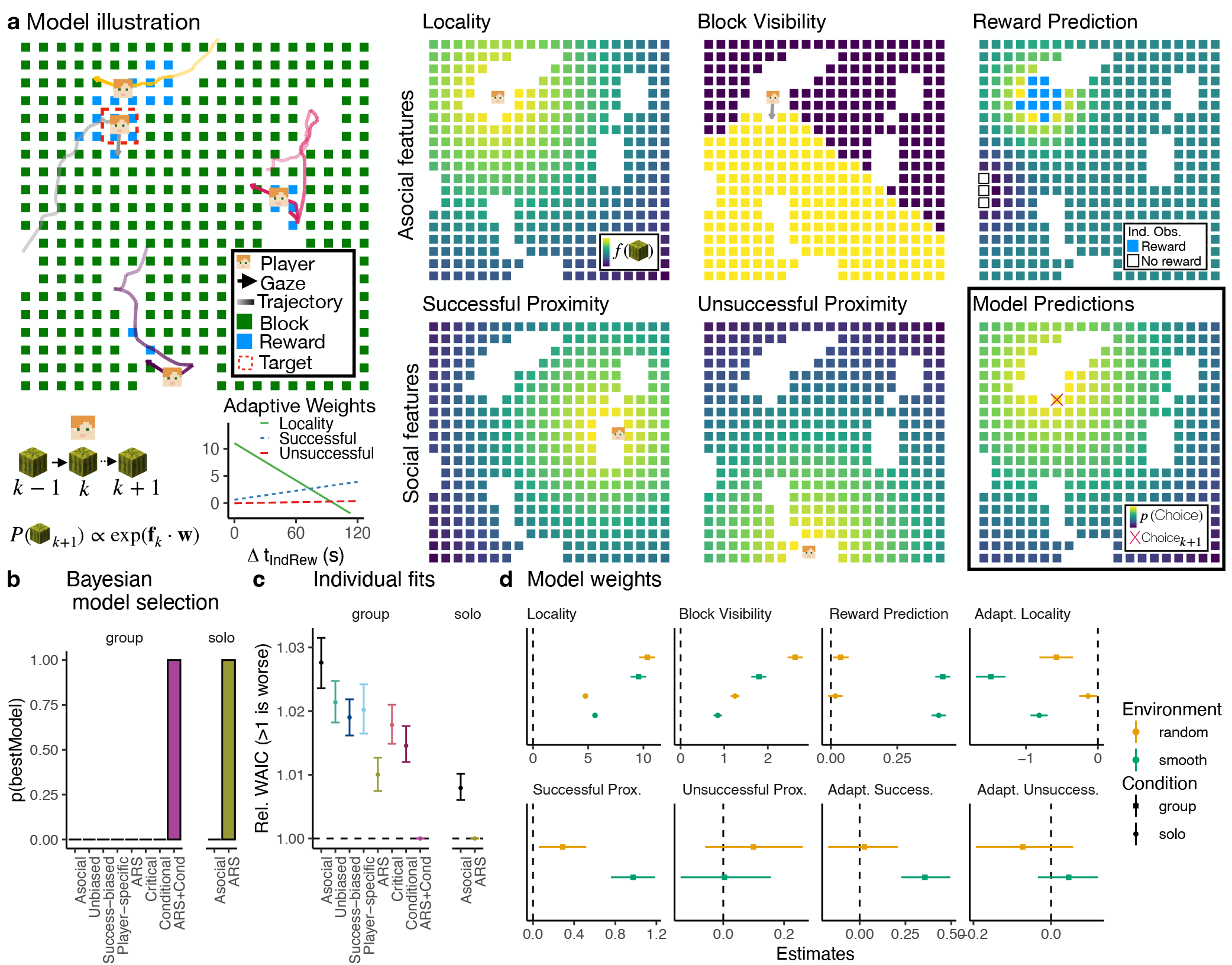
Computational model. (**a**) Left: Model illustration focusing on the player highlighted in the red dashed line. Right: Different models incorporate different sets of features (see text for details), where we illustrate the five features of the winning ARS+Cond model. Increasingly yellow colors correspond to higher feature values, with model predictions (bottom right) shown using the posterior mean of population-level weight estimates, with the red cross indicating the actual choice. ARS+Cond is a dynamic model, where locality and successful/unsuccessful proximity weights (bottom left) change as a function of time since the last individual reward. (**b**) Model comparison using protected exceedance probability^44^ to describe the posterior probability of the best model. (**c**) Individual model fits showing relative WAICs to the winning model (in each condition), with dots showing the group means and error bars indicating the 95% CIs. (**d**) Population-level weight estimates of the best models (group:ARS+cond; solo: ARS; see Figs. S12-S13 for all models), with the dots showing the posterior mean and error bars showing the 95% HPDI, and a vertical dashed line at 0.

### Behavioral results

To understand how the environment and social interactions impacted performance, we examine the normalized reward rate (Fig. 2a), which controls for faster reward depletion in group rounds with more individuals searching for the same number of finite rewards (see Fig. S1). Using a hierarchical Bayesian regression (see Methods), we find that participants acquired greater rewards in smooth environments (posterior mean: 0.22, 95% Highest Posterior Density Interval: [0.19, 0.24]), but with no reliable influence of social condition (0.002 [-0.02, 0.02]), nor interaction between condition and environment (−0.02 [-0.05, 0.01]). Thus, both individuals and groups achieved higher foraging success in smooth environments, whereas the simultaneous informational benefits and competitive drawbacks of social foraging did not yield any reliable changes to performance at the aggregate level. We also found no reliable increases in foraging success over rounds (random: 0.00 [-0.01, 0.02]; smooth: 0.01 [-0.01, 0.04]), and thus have no evidence of meta-learning during the task.

Yet, individual performance could be improved through adaptive foraging. To test this hypothesis, we first measured how foraging distance changes depending on whether the previous block yielded a reward (Fig. 2b). In smooth environments, participants foraged more locally after acquiring a reward (−1.6 [-2.0, -1.3]; no reliable effect of group, see Fig. S2a). In contrast, this effect was reversed in random environments, where participants foraged over larger distances after a reward (0.3 [0.1, 0.6]; no effect of group). In turn, greater adaptivity (i.e., difference in foraging distance) predicted better individual performance (Fig. S3) in smooth (solo: *r*_*τ*_ *=*.20, *p <* .001, *BF=* 32; group: *r*_*τ*_ .18, *p =* .003, *BF=* 9.9; Kendall’s tau) but not random environments (all *p >* .05; *BF <* 1). We obtained equivalent results when measuring the turning angle between blocks (Figs. S2-S4), consistent with the theoretical mechanisms of ARS^15^.

Next, as an initial analysis of social interactions, we computed the average pairwise distance between participants across conditions (Fig 2c). In solo rounds, the four participants foraged on separate but identical fields, allowing us to calculate an asocial baseline by superimposing their spatial coordinates as if they were on the same field. This analysis revealed greater social distancing in random compared to smooth environments (0.6 [0.8,0.4]; Fig. S5a), which was reliably larger when comparing group rounds to the asocial baseline (0.3 [0.1, 0.5]), suggesting an active increase in avoidance. In smooth environments, where participants generally foraged in greater proximity to each other, social distance was not reliably different from the asocial baseline (0.004 [-0.2,0.2]). Later, we show that this lack of difference in social distance at the aggregate level hides context-dependent changes (see Temporal dynamics).

Lastly, we used visual field transcription (see Methods) to measure social visibility between each pair of participants at a given time point. We first analyze the average number of visible peers (Fig. 2d), again computing an asocial baseline from solo rounds as if they were on the same field. In general, participants observed one another more in smooth environments (0.01 [0.001, 0.02]; Fig. S5b), although there was no reliable difference when comparing group rounds to the solo baseline (−0.0001 [-0.01, 0.01]). In contrast, participants reliably reduced social visibility in random environments compared to the asocial baseline (−0.01 [-0.02, -0.00005]), again showing active avoidance. Note that when comparing how visibility changes across conditions at the aggregate level, inbound and outbound social visibility are equivalent, masking individual differences and context-dependent adaptation. We now turn to network analyses to provide a better understanding of the structure and asymmetries of social interactions through visibility.

#### Proximity and visibility networks

Next, we performed social network analyses on spatial and visual interactions (Fig. 2e). *Proximity networks* describe each participant as a node with undirected edges weighted by the average proximity (i.e., inverse distance) between players. *Visibility networks* were constructed similarly, but with directed edges weighted proportional to the duration of time each target player was visible to another player. Network weights are computed per individual per round, and then averaged per individual (within each condition), with the same analyses also applied to solo rounds to provide an asocial baseline.

We first used the proximity network to compute the Eigenvector centrality for each participant, providing a holistic measure of the influence each node exerts on the network. Higher centrality corresponds to participants who maintain close proximity to others, especially to those with high proximity scores themselves. Whereas centrality did not predict rewards in random environments (group: -7.6 [-16.6,0.6]; solo: -7.6 [-16.1,0.8]; all slopes overlap with 0), we found a robust inversion in smooth environments, where higher centrality predicted higher rewards in group rounds (7.9 [1.6,14.0]; Fig. 2f). This effect disappeared in the asocial baseline (2.6 [-5.6,10.1]), suggesting that the benefits of spatial centrality were due to social dynamics. However, it is unclear whether being central facilitated better performance (via access to social information) or if centrality resulted from better performance (via effective success-biased imitation), motivating the need for dynamic analyses (see below).

Next, we examined the relationship between in- and out-degree in the visibility network. In-degree is the sum of all inbound edge weights, where being observed by more peers and for longer durations both contribute to larger in-degrees. Similarly, higher out-degree corresponds to observing other peers with longer durations. This analysis revealed an asymmetry in social attention, with a general inverse relationship between in- and out-degree (Fig. 2g). Whereas this asymmetry was also present in random and solo rounds, it was reliably stronger when combining group rounds and smooth environments (group+smooth interaction: -0.49 [-0.89, -0.08]). This suggests an increased specialization of social learning strategies (i.e., “producers” with low out-degree and high in-degree or “scroungers” with high out-degree and low in-degree) and asymmetry of social attention in settings where social information was useful. Thus, our network analyses provide insights into asymmetric patterns that could not be detected at the aggregate-level (Fig. 2d). However, neither in- or out-degree predicted individual performance (Fig. S10). We now analyze temporal dynamics to better understand how social interactions change over time and in response to different contextual factors.

#### Temporal dynamics

To analyze the dynamics of social interactions, we searched for temporally predictive clusters relating individual reward to our high-resolution spatial-visual data (Fig. 2h-k; see Methods). More precisely, we computed correlations between time-series at different temporal offsets (Fig. 2h), and searched for temporally continuous clusters of significance (indicated by bold lines in Fig. 2i-k) that survived multiple forms of chance correction (see Methods). At negative offsets, these clusters indicate that reward predicts future proximity/visibility (either positive or negative correlations depending on the sign), while positive offsets indicate that proximity/visibility predicts future reward.

First, the dynamics of individual reward (i.e., foraging success) and spatial proximity to other players revealed a pattern of *success-dependent spatial cycling* in smooth (but not random) environments (Fig. 2i). The positive correlation at offset -19s to -2s (bold line) indicates that greater reward predicted greater proximity at long timescales into the future. Since previous analyses showed that success corresponded to reduced foraging distances (Fig. 2b) and increased turning angles (Fig. S2b), the greater proximity predicted by reward is more likely due to attracting peers when successful. Subsequently, this creates a cyclical pattern, where the negative correlation at offset 9s to 20s indicates that greater proximity predicts lower future rewards, likely due to depletion. Lower rewards then drives reduced proximity (i.e., reward → proximity effect at negative offsets), with reduced proximity driving increased rewards, thus continuing the cycle. Crucially, this pattern is absent in random environments (flat effect) and qualitatively different in the asocial baseline (unimodal positive correlations centered at 0s; see Fig. S7), allowing us to rule out the role of the smooth resource distributions. These dynamics reveal the mechanisms behind why greater centrality was correlated with performance (Fig. 2f), showing that success temporarily attracts imitators, who then dissipate when resources are depleted.

Next, we looked at the dynamics of reward and visual field data, where we analyzed the number of visible peers (out-bound social attention towards others; Fig. 2j) and the number of observers (inbound social attention; Fig. 2k) at every time-point. Starting with outbound visibility of peers (Fig. 2j), we found evidence for *adaptive social attention*. First, we found several negative correlation clusters at negative offsets for both environments, indicating that low rewards predicted greater visual attention towards others (and vice versa). However, only the cluster in smooth environments survives (from -2.85s to -0.45s) when computing the contrast between group and solo rounds (Fig. S7). Thus, our comparison to the asocial baseline allows us to identify that social attention adapts to individual performance, above and beyond the generic effects of the task structure (i.e., reduced visibility when destroying a block). Additionally, we also observe opportunity costs of social attention in smooth rounds (1.3s to 2.3s), where the negative correlation indicates that more visible peers predicted lower rewards. This effect is absent in random environments and reversed in the asocial baseline (Fig. S7), which accounts for the shared reward structure, indicating it is a distinctly social phenomenon.

Lastly, the dynamics of reward and inbound visibility (number of observers) indicate *success-biased selectivity*, where participants who acquired higher rewards were the target of social attention (Fig. 2i). In smooth environments, we observed two clusters at negative offsets (−13.6s to -3.7s and -3.0s to - 1.0s) with positive correlations, indicating that greater rewards predicted more observers in the future. This success-biased selectivity was absent in random environments and inverted in the asocial baseline (Fig. S7). The strong dip approaching offset = 0s is due to the visual dynamics of the task, since the splash animation temporarily obscured the avatar when acquiring a reward. Additionally, we observe another positive correlation cluster at offsets 1.0s to 2.2s in smooth environments, indicating having more observers predicted greater future rewards. This effect was again absent in random environments and inverted in the asocial baseline (Fig. S7).

In sum, these temporal dynamics reveal how individual performance is the key driver of adaptive mechanisms, driving changes in social proximity (Fig. 2i) and social attention— both when (Fig. 2j) and towards whom (Fig. 2k) it is directed. We next relate these social attention processes to social interactions characterized by concrete leader-follower dynamics.

#### Social influence and leadership

Inspired by methods used to study collective decision-making in wild baboons^42^, we analyzed the frequency of “pull” events that capture leader-follower dynamics (see Methods). Each candidate event was selected from min-max-min sequences in the pairwise distance between players (Fig. 3a) and then filtered by a number of criteria including strength (change in distance relative to absolute distance) and disparity (one player moves more than the other). After filtering, we detected a total of 537 pull events (see Fig. 3a for an example), where in each event, one player is identified as a *leader* (moved more during [*t*_1_, *t*_2_]) and the other as a *follower* (moved more during [*t*_2_, *t*_3_]).

We analyzed both solo and group rounds, with solo rounds providing a benchmark for the sensitivity of these analyses by accounting for the influence of the reward structure (Fig. 3b). While random environments saw a reduction in pull events from solo to group rounds (hierarchical Poisson regression: -0.7 [-1.2, -0.1]), smooth environments saw a large increase in pull events from solo to group rounds (1.4 [0.8, 2.0]). These results were robust to different filter thresholds (Fig. S8) and suggest participants not only adapted their social attention (Fig. 2g,j) but also their susceptibility to social influence depending on the relevance of social learning: following others when adaptive (smooth), and actively avoiding others when maladaptive (random).

Next, we computed a leadership index for each participant based on their frequency of being a leader vs. a follower: *n*_leader_ − *n*_follower_, using only smooth group rounds for interpretability. Participants with a high leadership index were observed more (i.e., higher in-degree) and observed others less (i.e., lower out-degree), indicating a high correspondence between our analysis of these non-overlapping aspects of the data (i.e., visual field data vs. spatial trajectories; see Fig. S9). Yet neither leadership (Fig. 3c) nor in/out-degree predicted performance (Fig. S10f). However, when we focus on the instantaneous reward rate *during* a pull event (Fig. 3d), we found that leaders received more rewards than followers (0.6 [0.3,0.9]). Thus, social influence appears to be modulated by success bias, although we find no long-term benefits of social attention or leadership at the behavioral level, motivating the need for more precise computational modeling (see below).

#### Behavioral summary

Overall, participants *adaptively* deployed both asocial and social learning mechanisms according to the environment and depending on individual performance, and *selectively* directed social attention towards successful individuals. These behavioral results provide a lens into the dynamics of how asocial and social learning interact and feedback onto each other. However, they only indirectly speak to the individual-level cognitive processes that drive decision making in our experiment. Therefore, we next turn to computational models integrating different combinations of asocial and social learning mechanisms to predict individual foraging decisions.

### Computational modeling of choices

We use a computational modeling framework (Fig. 4a) to sequentially predict which block participants will destroy next:

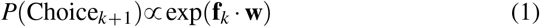

Predictions are modeled as a softmax distribution over a linear combination of block features **f** and weights **w**, where we use the state of the world when the *k*-th block is destroyed in order to predict the *k +* 1-th block. Block features **f** capture hypotheses about individual and social learning mechanisms (see below), while weights **w** are estimated using hierarchical Bayesian methods, accounting for individual and group variability as random effects (see Methods).

#### Asocial features

We first used a set of asocial features to capture physical constraints of the task and individual learning through reward prediction (Fig. 4a; see Methods). *Locality* is the inverse distance to the player at time *k*, reflecting a tendency to forage locally. *Block Visibility* captures which blocks are within the player’s field of view at time *k*, and is set to 1 if visible and 0 if not. *Reward Prediction* uses Gaussian Process regression as a cognitive model of asocial reward generalization in structured environments^37,43^. Since each block can only be destroyed once in each round, reward prediction relies on value function approximation, as a common form of generalization in reinforcement learning^18^, where past observations are used to infer a value function over the search space. Here, we implement this as a binary classification problem, where based on the player’s reward history (until time *k*), we predict the probability of each remaining block containing a reward as a logistic sigmoid of an inferred latent variable *z* (Fig. 4a Reward Prediction panel), with higher values corresponding to higher probability of reward (see Methods).

#### Social features

We then incorporate social features based on proximity to different subsets of players, motivated by our results showing selective visual attention (Fig. 2j) and a susceptibility to being “pulled” (Fig. 3d) towards successful individuals. *Successful Proximity* is computed using players who were visible and were observed acquiring a reward (i.e., visible splash) in the span of *k −* 1 to *k* (using visual field transcription). We used the last observed location of each player to compute proximity (inverse distance), and use the centroid if there were multiple successful players. *Unsuccessful Proximity* is calculated the same way, but for visible players who were not observed acquiring a reward. In separate models, we also tested *Social Proximity* to all players (irrespective of success) and *Player-specific Proximity*, with unique weights for each target player (see Methods).

#### Model comparison

We compared a series of models, each using a different subset of features. The models fall under one of two classes: *static models* in which the weights remain constant over a round (i.e,. fixed strategy), and *dynamic models* in which the weights adaptively change (Fig. 4a bottom left) as a function of the elapsed time Δ*t* since the last individual or socially observed reward (depending on the model):

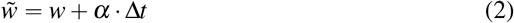

In dynamic models, the weight estimate *w* acts as an intercept, while *α* captures the degree of adaptivity as a slope. Since we first normalize Δ*t*, weight estimates are comparable between static and dynamic models. Model comparison is performed separately for group and solo rounds, where we use Bayesian model selection^44^ to compute the probability of the best model (i.e., protected exceedance probability; Fig. 4b) and also report individual WAICs relative to the best model (Fig. 4c) for more fine-grained analysis.

Our static models include 1) an *Asocial* model using only asocial features, 2) an *Unbiased* model that adds undifferentiated Social Proximity across all players as a naïve form of social imitation, 3) a *Success-biased* model using separate weights for Successful vs. Unsuccessful players, and 4) a *Player-specific* model with separate proximity weights for each player. Of these models, the Success-biased model performed best in group rounds (Fig. 4c).

Our dynamic models are inspired by influential theories of adaptivity in asocial and social foraging. 5) *ARS* is based on past work using area-restricted search^15,33^, and uses only asocial features, but with the locality weight changing as a function of time since the last individual reward. This corresponds to the common finding^40^ that search distance adapts to foraging success (Fig. 2b), and was the best model in solo rounds and the second best in group rounds (Fig. 4b-c). We also developed two adaptive social learning models inspired by Enquist et al.^4^, both based on our success-biased model. 6) The *Critical learner* adapts the reward prediction weight as a function of time since the last socially observed reward, while 7) the *Conditional learner* adapts both the successful and unsuccessful proximity weights as a function of time since the last individual reward. Thus, the critical learner adapts individual learning as a function of social learning success, while the conditional learner adapts social learning as a function of individual success, with the latter performing better (Fig. 4c). Finally, we combined ARS with the Conditional learner to create 8) an *ARS+Cond* hybrid, where individual performance drives adaptivity of both asocial (i.e., locality weight) and social (i.e., success-biased imitation) mechanisms. This ARS+Cond model vastly outperformed all other models in group rounds (*p* bestModel > .999; Fig. 4b) and was better than the sum of its parts (i.e., ARS or Conditional learning alone; Fig. 4c).

#### Model weights

We focus on interpreting the weights for the best models in each condition (group: ARS+Cond; solo: ARS), but all models had similar weights for shared features (Figs. S12-S13).

Locality and Block Visibility influenced choices in all conditions, and were typically stronger in random than in smooth environments (i.e., in the absence of reward-predictive cues). The one exception is that in solo rounds, participants foraged more locally in smooth than random environments (0.9 [0.8, 0.9]), suggesting greater local exploitation in the absence of social competition. Reward Prediction weights were reliably larger in smooth environments (0.41 [0.38, 0.45]; no reliable difference between group-solo: 0.02 [-0.02, 0.06]), but overlapped with 0 (solo: 0.02 [-0.01, 0.04]) or were negligible (group: 0.04 [0.01, 0.07]) in random environments. Thus, participants adapted individual reward prediction based on the environment, and this mechanism was unaffected by the social context. Notably, we found adaptation in locality weights in all conditions (i.e. *ARS*), with stronger adaptivity in smooth environments (group:-0.91 [-1.05, -0.77]; solo: -0.68 [-0.76,-0.60]) and in the group condition (−0.56 [-0.87, -0.23]). The negative sign corresponds to a reduction of locality as the participant experienced longer durations without individual rewards, consistent with past models of asocial foraging^15^. Our results expand on these previous findings, by showing that adaptation of local search increases (rather than diminishes) in social settings.

Social feature weights (ARS+Cond) show that participants were strongly influenced by successful players in smooth environments (1.0 [0.8, 1.2]), and substantially less in random environments (smooth - random: 0.7 [0.5, 0.9]). However, even in random environments the effect was reliably different from chance (0.29 [0.07, 0.53]), suggesting a persistence of success-biased imitation even in environments where social learning was irrelevant. In contrast, we found no effect of unsuccessful players in neither smooth (0.004 [-0.14, 0.15]) nor random environments (0.10 [-0.07, 0.25]). We also only observe adaptivity in the successful proximity weights for smooth (0.4 [0.2, 0.5]) but not random environments (0.10 [-0.07, 0.25]). Thus, participants increased their reliance on social information as they experienced longer periods without individual reward, with increased selectivity towards successful players.

Altogether, our modeling framework allows us to integrate theories of asocial and social adaptation from past literature^4,15^, made possible by our combination of visual and spatial data. Here, we show that these adapative mechanisms are driven by individual performance, and themselves adapt to the context of different reward environments. Furthermore, we can link individual weight and adaptability estimates to a number of behavioral signatures such as performance, centrality, and visibility (see Fig. S14). Notably, greater social adaptation predicted better performance in group rounds (Adapt. Success: 2.99 [0.41, 5.58]; smooth rounds), while greater asocial adaptation weights predicted better performance in solo rounds (Adapt. Prox: -2.34 [-3.72,-1.03]; smooth rounds). Thus adaptivity of both social and asocial learning mechanisms is what predicts individual performance.

## Discussion

Collective foraging is a common metaphor for human social learning^40,41,45^. With similar dynamics as marketplace innovation or scientific research, peers can be both useful sources of social information, but also competitors for the same limited resources. Here, we used an immersive virtual environment (Fig. 1), where each individual’s limited field of view imposes a trade-off between allocating attention to individual or social learning, while spatial proximity to others directly shapes opportunities (and also costs) for social interactions. With access to visual field data (Fig. 1c) and spatial trajectories, our analyses provide insights into the structural and temporal dynamics of social interactions, where we study how people adapt to both different reward environments (smooth vs. random; Fig. 1e) and to dynamically changing social environments.

Our results shed light on the adaptive mechanisms driving collective human behavior, integrating past theories from asocial foraging^15^ with context-dependent^4,9,10^ and selective^5,45,46^ mechanisms of social learning, which have yet to be combined in a single framework. This integration reveals how these asocial and social mechanisms amplify one another, and are driven by a common currency of individual performance, instead of being governed by separate cues that mutually compensate for one another. Furthermore, it is the degree of adaptivity—for both social and asocial learning mechanisms—that best predicts individual performance.

When rewards were smoothly clustered (offering traction for social learning), participants specialized more strongly with greater asymmetry of social attention (Fig. 2g), adaptively sought out social information depending on performance (Fig. 2j), and selectively directed their social learning towards successful individuals (Fig. 2k). Participants were also more likely to be “pulled” into leader-follower dynamics in smooth environments (Fig. 3b), which were selectively directed towards “leaders” with higher instantaneous reward rates (Fig. 3d). Our computational models (Fig. 4) combined spatial and visibility data to account for both asocial and social learning mechanisms, which dynamically adapt over time. Here, the winning model combined area-restricted search^15^ with conditional social learning^4^ (ARS+Cond), where individual performance was the key factor driving adaptivity of both local foraging and selective social learning (Fig. 4d). Notably, individual estimates of adaptivity of both asocial (i.e., ARS) and social learning mechanisms (i.e., success-biased imitation) predicted better individual performance (Figs. S3,S4, & S14). Overall, this work integrates previously disparate theories of adaptive mechanisms of asocial and social learning, providing new evidence that they are driven by a common mechanism of individual performance, with their interaction yielding an amplification of adaptivity and selectivity.

A long tradition of research has successfully generalized findings from spatial foraging to more abstract and general settings, such as searching for information on the internet^47^ or recovering internal memories^48^. Since then, recent advances in animal research have incorporated rich behavioral data from visual field analysis^49^, spatial trajectories^50^, and network dynamics^51^ with important new implications, showing how simple (and sometimes seemingly maladaptive) social learning mechanisms can give rise to intelligent behavior in dynamic and ecological environments. Yet, prior to this work, we have lacked behavioral data from humans in an equally immersive setting, which would allow us to account for similar mechanisms. Here, our immersive Minecraft setting has allowed us to address this gap, linking asocial and social mechanisms of adaptive behavior with potential future implications for human social learning^52^.

### Limitations and future directions

While substantially reduced, success-biased social learning was also present in random environments (Fig. 4d). Thus, despite resources being distributed randomly, participants were still somewhat drawn towards successful peers. One explanation is that success-biased copying is hard for people to unlearn (e.g., similar to imitation^53,54^), since it is beneficial in many settings. Alternatively, participants may have believed there to be structure in random environments, as is often the case when observing random events (e.g., the gambler’s fallacy^55^). Either way, these results suggest potential limitations to the degree of human adaptability and a lingering bias towards social learning. However, even though social information provided no benefits in random environments, it may still offer a computationally cheap tool for engaging in exploration^11^ (away from one’s current location). Individual exploration is associated with cognitive costs and can be impaired by imposing memory load^56^ or time pressure^57^. Thus, social imitation may act as an “exploration device” at a reduced cognitive cost relative to individual exploration^11,38,58^. An extreme test could be to use anti-correlated environments to observe if success-biased social learning still persists in the most non-adaptive settings. Indeed, different environments could also yield greater evidence for critical social learning over conditional social learning^4^, where costlier asocial learning may prioritize using social cues to adapt learning mechanisms.

The asymmetry of social attention (amplified in smooth environments; Fig. 2g) may act as a safeguard against maladaptive herding^12,59^, where instead of copiers copying other copiers, social learning is selectively directed towards individual learners (low out-degree) and is selectively directed towards successful individuals (Fig. 2k). In our study, successful foraging outcomes were made salient by a visual cue (i.e., splash), although people can also deploy metacognitive strategies to infer latent performance or skill from overt behavior^5,14^, providing additional mechanisms for guiding selective social learning. Future work can explore the extent to which these mechanisms (together with our ability to discount correlated social information^60^) may offer a degree of natural protection against the spread of misinformation^61^ and the formation of echo chambers through homophilic social transmission^62^.

Future work may consider using a non-depleting reward environment, where collective coordination can yield additive benefits to individual search^5,63–65^. Indeed, a better understanding of our ability to cumulatively innovate upon previous solutions over long multi-generational timescales has been a powerful motivating force in social learning research^2,27,66^. Here, we have focused on understanding the temporal dynamics of social learning over short timescales, which produced novel insights into the cognitive mechanisms supporting flexible and adaptive social learning. However, a more complete understanding requires connecting social learning mechanisms observed at short timescales to adaptive outcomes over long, cultural timescales. Our work provides the foundations for this endeavor, by providing insights into the cognitive mechanisms that make people such powerful social learners in dynamic and more realistic contexts.

Our task was originally designed for data collection using a virtual reality (VR) headset instead of a mouse and keyboard. However, preliminary testing revealed that locomotion via teleportation (the preferred method to avoid VR motion sickness) resulted in less naturalistic spatial trajectories and interfered with visual field analyses due to the field of view temporarily fading to black during movement. Several VR treadmill products were also tested as an alternative form of locomotion, but required substantial training and resulted in large individual differences in navigation ability. In contrast, the current computer-based modality captured naturalistic trajectories in both space and heading direction. However, future studies should continue to strive for greater ecological validity using new advances in VR.

In conclusion, our study of collective foraging in an immersive Minecraft environment integrated computer-transcribed visual field data with high-resolution spatial trajectories to provide a integrative perspective on the adaptive mechanisms of asocial and social learning. Ultimately, this work advances our understanding of the cognitive mechanisms underlying adaptive learning and decision making in social contexts, and provides the foundation for future investigations in non-spatial domains of social interactions.

## Methods

### Participants and design

This research complies with all relevant ethical regulations and was approved by the Institutional Review Board of the Max Planck Institute for Human Development (MPIB; approval number: A 2019-05). Participants (*n =* 128) were recruited from the MPIB recruitment pool in Berlin (82 female; *M*_age_ =27.4, *SD*_age_ =5.0) and participants signed an informed consent form prior to participation. No statistical method was used to predetermine sample size and no data were excluded from the analyses. Participants received a base payment of € 12 plus a bonus of € 0.03 per reward, spending approximately one hour and earning on average € 17.32 ± 1.02 (*SD*).

Participants completed the task in groups of four. After an in-game tutorial (Supplementary Video 1) and two practice rounds (see below), participants completed 16 2-minute rounds of the task. Using a within-subject design, we manipulated the reward structure (random vs. smooth; Fig. S15) and search condition (solo vs. group). The order of round types was randomly assigned and counterbalanced across groups, with four consecutive rounds of the same type (Fig. 1d). The study coordinator was blinded to the allocation during the experiments. The reward structure and search condition for each round was announced prior to the start of each round in an onscreen notification.

The reward structure of a given round was made salient by mapping each reward structure to either pumpkin or watermelon blocks (counterbalanced across groups). In both reward structures, 25% of blocks contained rewards, but rewards were either randomly or smoothly distributed. The smooth environments were generated by sampling from a Gaussian process^67^ prior, where we used a radial basis function kernel (Eq. 12) with the lengthscale parameter set to 4 (similar to^37^). Sampled reward functions were then binarized, such that the top quartile (25%) of block locations were set to contain rewards. We generated 20 environments for both smooth and random conditions (Fig. S15), with each session (i.e., group) sampling without replacement 1 (practice) + 8 (main task) = 9 environments of each class with pseudorandom assignments that were pregenerated prior to the experiment. In the tutorial (Fig. 1d), participants were given verbal descriptions of each reward condition, saw two fully revealed illustrations of each environment class from a bird’s-eye perspective, and interactively destroyed a 3×3 patch of both smooth and random environments (Supplementary Video 1).

The search conditions were made salient by having participants stand on a teleportation block either by themselves (solo) or with the other three participants (group) in order to begin the round. In the solo condition, participants searched on identical replications of the same environments but without interacting with each other. In the group condition, participants searched on the same environment and could interact with, compete against, and imitate one another.

### Materials and procedure

The experiment was implemented as a computer-based experiment, with each computer connected to a modified Minecraft server (Java edition v.1.12.2). In the experiment, the sound was turned off, participants could not see each other’s screens, and task-irrelevant controls (e.g., jumping, sprinting, inventory, etc) were made unavailable. The Minecraft world consists of “blocks” that can be “mined” for resources by holding down the left mouse button to hit them until they are destroyed. In the experiment, participants controlled an avatar that moved through our custom-made environment, defined as a flat field containing a 20×20 grid of 400 pumpkin or watermelon blocks (Fig. 1a) with a two block space between each block. The field was bounded by an impassable fence. See Supplementary Video 2 for a bird’s-eye illustration of a round, and Supplementary Videos 3 and 4 for screen captures from group rounds on smooth and random reward environments, respectively.

Each resource block (either watermelon or pumpkin) could be foraged by continually hitting it for 2.25 seconds until it was destroyed, yielding a binary outcome of either reward or no reward. Rewards were indicated by a blue splash effect, visible by other players from any position if it was in their field of view. Only resource blocks could be destroyed in the experiment and destroyed block were not renewed. Blocks did not possess any visual features indicating whether or not they contained a reward. However, rewards in smooth environments were better predictable, since observing a reward predicted other rewards nearby. Participants were individually incentivized to collect as many rewards as possible, which were translated into a bonus payment at the end of the experiment. The cumulative number of rewards (reset after the practice rounds) was shown at the bottom of the screen.

After receiving verbal instructions, participants completed an in-game tutorial to familiarize themselves with the controls, how to destroy blocks, the difference between smooth and random reward distributions, and the overall task structure (Supplementary Video 1). They then completed one solo practice round in a smooth environment and one solo practice round in a random environment. These were identical to the solo condition of the main task, but performance in these rounds did not contribute to a participant’s bonus payment. Each round lasted 120 seconds, with the end of the round corresponding to the sun setting below the horizon. This served as an approximate in-game timer for each round, and was communicated to participants in the tutorial. A 3-second countdown timer was also shown onscreen. At the end of the round, participants were given an onscreen announcement indicating the number of rewards they had earned and notifying them of the reward structure and search condition for the next round. Participants were then teleported into a lobby (separate lobbies for solo rounds or a communal one for group rounds), and were required to all stand on a “teleportation” block to indicate readiness for the subsequent round. Prior to the start of a social round, participants all stood on a communal teleportation block, while prior to solo rounds, participants each stood on separate teleportation blocks, in order to induce the social or asocial context. Once all players were ready, a 3-second countdown was displayed and they were teleported into a random position in the next environment with a random orientation direction.

## Data collection

Experimental data was collected using a custom data logging module programmed in Java, which were separated into map logs and player logs. Map logs recorded information about each block destruction event, including a timestamp, player identifier, block position, and the presence or absence of reward. Player logs contained each player’s position in the horizontal XZ-plane together with the XYZ components of their heading vector (i.e., where they were looking). Both logs contained information sampled at Minecraft’s native 20 Hz tick-rate (i.e., once every 0.05s), providing high-resolution data about spatial trajectories and heading directions.

## Automated transcription of visual field data

We developed a custom tool built on the Unity game engine (ver. 2019.3) for performing automated transcription of visual field data (Fig. 1c). We first used data collected from the experiments to simulate each round of the experiment from each participant’s point of view. These simulations were then used to automate the transcription of each participant’s field of view (Supplementary Video 5).

Our Unity simulations assigned each entity in the experiment (i.e., each block, player, and reward event) a unique RGB value, which was drawn onto a render texture one tenth the size of the player’s actual monitor (192×108 pixels as opposed to 1920×1080 pixels). Since the images were rendered without any anti-aliasing or transparency through a simple, unlit color shader, the RGB value of any drawn pixel could be uniquely related with a lookup table to the corresponding entity. We then simulated each round of all experiment data from each player’s perspectives within the Unity game engine, using the map logs and player logs, which allowed us to fully reconstruct the world state. Once all four player perspectives were individually rendered, we could read out the pixels from each player’s field of view, using the RGB colors of the simulated pixels to determine whether an entity was visible at any point in time (20 Hz resolution), and what proportion of the screen it occupied.

In creating these simulations, a few approximations were required. In addition to the reduced resolution mentioned above, player models were approximated by their directionally-oriented bounding box and we ignored occlusion from the heads up display and view-model (e.g., occlusion due to hand position of the avatar). Additionally, some animations produced by the Minecraft game engine include inherent stochastic processes that were approximated. Namely, the splash particles used to indicate a reward event are generated in Minecraft using a random process that spawns 300 particles at predefined locations in a sphere around the player. Whilst the starting locations are deterministic, small deviations in velocity and the lifetime of these particles are generated randomly. Thus, we tuned the parameters of Unity’s particle system to be as authentic as possible by comparing simulated splash effects with video footage of splash effects generated by the Minecraft game engine.

We used a similar procedure for the solo rounds to establish an asocial baseline for our analyses. Whereas all four players searched on different replications of the same field, we simulated the solo rounds as if they were superimposed on the same field. Again, a few approximations were required. In these solo simulations, we removed a block whenever any of the four players destroyed it. Additionally, we generated a splash for each reward event, meaning if multiple players foraged the same block in a round, it would trigger a different splash event each time.

### Agent-based simulations

We implemented agent-based simulations to understand how the different reward environments (smooth vs. random rewards) interact with individual-level learning strategies (asocial learning vs. unbiased imitation vs. biased imitation) in determining foraging success (see Fig. 1f). The simulations uses the same features as the computational model, but are defined for a simplified version of the task, capturing the key visual-spatial dynamics of collective decision-making.

More precisely, our simulations modeled the foraging task as a discrete-time sequential game with partial observability, which generalizes Markov decision processes to incorporate multiple agents, partial observability, and separate rewards^68^. Formally, a task is a tuple, ⟨ 𝕀, 𝕊, 𝔸, 𝕆, *T, R, O⟩* : an agent index set, 𝕀; a set of environment states corresponding to configurations of agent locations/directions and available/destroyed blocks, 𝕊; a set of joint actions corresponding to agents moving in cardinal directions, 𝔸= × _*i*_ 𝔸_*i*_; a set of joint observations, 𝕆 = × _*i*_ 𝕆_*i*_, where each 𝕆_*i*_ is a subset of events viewed from agent *i*’s perspective (i.e., other agents’ locations, reward reveal events, and available blocks); a environment transition function, *T* : 𝕊 × 𝔸 → 𝕊; a joint reward function *R* : 𝕊×𝔸×𝕊 → ℝ^|𝕀|^; and a joint observation function, *O* : 𝔸 ×𝕊 →𝕆.

Agents are modeled as selecting a destination to navigate to, navigating to that destination, and then destroying the target block (requiring *k =* 9 timesteps in the simulation; approximately equivalent to the 2.25 seconds required to destroy a block and the maximum movement speed of 4.3 blocks/second). Agent policies consist of a high-level controller that transitions among different modes of behavior *n* ∈ {`SelectDest`, `NavTo (`*x*), and `forage (`*k*)}, where *x* is a target destination that a low-level navigation controller moves towards and *k* is a counter for the number of timesteps left to complete the destruction of a block. When in `SelectDest`, the controller samples a destination from *P*_**w**_(*x*)*∝*exp {**f (***x*) · **w**}, where **f** : *X →* ℝ^*K*^ returns a real-valued feature vector (incorporating both asocial and social mechanisms, the same as in the computational models; see below) for each destination block, and **w** ∈ ℝ^*K*^ are feature weights.

We considered populations of three types of agents. *Asocial agents* used a combination of locality (distance from current location), block visibility (using a 108.5-degree field of view as in the experiment), and asocial reward learning (see the subsection “Gaussian process for binary reward prediction” below). *Unbiased social agents* added an additional feature using the average proximity from observed social partners since the last choice, while *biased social agents* used a similar social proximity feature, but computed only from social partners that were observed acquiring a reward since the last choice. All feature weights were arbitrarily set to 1. For each of 20 random/smooth environments, we generated 100 simulations for each agent type in groups of four agents (for a total of 20 × 2× 100 ×3 = 12, 000 simulations). Each simulation was run for 400 timesteps. Figure 1f provides the results of the simulations, showing the average total reward collected by agents by environment type (smooth/random) and strategy (asocial/unbiased social/biased social).

### Hierarchical Bayesian regressions

Statistical analyses were conducted using hierarchical Bayesian regressions to simultaneously model the effects of the experimental manipulations (smooth vs. random and solo vs. group), while accounting for participant- and group-level variability. All regression models used Hamiltonian Markov chain Monte Carlo (MCMC) with a No-U-Turn sampler^69^ and were implemented using `brms`^70^. For count-based variables (e.g., blocks destroyed or pull events), we used Poisson regression, but report the un-transformed regression coefficients for simplicity. All models used generic, weakly informative priors ∼𝒩(0, 1) and all fixed effects also had corresponding random effects following a maximal random-effects procedure^71^. All models were estimated over four chains of 4,000 iterations, with a burn-in period of 1,000 samples. Effects are reported as the posterior mean and 95% Highest Posterior Density Interval (HPDI).

### Temporal dynamics

Based on methods developed in neuroscience^72^, the temporal dynamic analyses (Fig. 2h-k and Fig. S7) use time-series data from each participant in each round to discover temporal structures in social interactions, where rewards predict future spatial/visual patterns (negative offsets) and spatial/visual patterns predict future rewards (positive offsets).

The time-series variables we used are reward (binary vector), spatial proximity (average inverse distance to all other players), and both the number of visible peers and the number of observers (integer variables ∈ [0,3]; acquired from the automated transcription of visual field data) at every point in time (20 Hz time resolution). For solo rounds, we computed both spatial proximity and visibility as if participants were on the same field to provide an asocial baseline.

We then computed correlations between each pair of variables `cor(V1,V2)`, where we iteratively time-lagged `V2` from -20 to +20 seconds, with non-overlapping regions of each time series omitted from the data (see Fig. 2h). Each correlation was then *z*-transformed and corrected for chance using a permutation baseline (i.e., permutation chance correction). This chance correction is based on iteratively permuting the order of `V2` and computing the correlation `cor(V1,V2_permuted)` over 100 different permutations (for each correlation). We then subtracted the *z*-transformed mean of the permutation correlations from the target correlation. These permutation corrected correlations are reported as a population-level mean (± 95% CI) in Figure 2i-k and Figure S7.

Lastly, to provide better interpretability of these results, we used a maximum cluster mass statistic^72^ to discover temporally continuous clusters of significance at the population level. For each pair of variables [*V* 1,*V* 2] and within each combination of condition (solo vs. group) and environment (random vs. smooth), we used a cluster permutation test to find a threshold for random clusters. This analysis used 10,000 permutations, where for each, we iterated over each individual time series of *z*-transformed (and chance-corrected) correlations, randomly flipping the sign at each time point. We then used a single-sample *t*-test with *α* = .05 to compute which time points (at the population level) were significantly different from 0. This provided a distribution of the duration of temporally continuous clusters of significance in the randomly permuted data. We then used the upper 95% CI of this distribution as a minimum threshold for the actual data, where we applied the same significance testing procedure, but discarded all clusters shorter in duration than the permutation threshold. The surviving clusters are illustrated with bold lines in Figure 2i-k and Figure S7.

### Social influence

We used methods developed to analyze the movement patterns of geotracked baboons in the wild^42^ to measure social influence. This allows us to detect discrete “pull” events over arbitrary time scales, where the movement patterns of one participant (leader) pull in another (follower) to imitate and forage in the same vicinity (Fig. 3).

We first computed the pairwise distance between all participants (Fig. 3a) and defined candidate pull events from min-max-min sequences, where we used a noise threshold of 1 block distance to determine what corresponds to minimum and maximum distances. These candidate sequences were then filtered based on strength, disparity, leadership, and duration in order to be considered a successful pull.

**Strength** *S*_*i, j*_ defines the absolute change in dyadic distance relative to absolute distance:

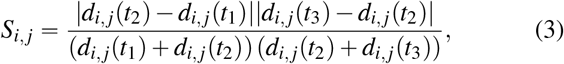

where *d*_*i, j*_ (*t*_*k*_)is the dyadic distance between participants *i* and *j* at time *k* [1, 2, 3] (corresponding to the timepoints of the min-max-min sequence). We required pull events to have a minimum strength of *S*_*i,> j*_ .1, such that they correspond to meaningful changes in spatial proximity rather than minor “jitters” at long distance.

**Disparity** *δ*_*i, j*_ defines the extent to which one participant moves more than the other in each segment, relative to the total distance moved by both participants:

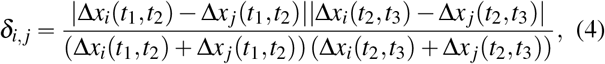

where Δ*x*_*i*_ (*t*_1_, *t*_2_)is the displacement between *t*_1_ and *t*_2_. We filtered pull events to have a minimum disparity of *δ*_*i, j*_ *>* .1, such that changes in spatial proximity were asymmetrically driven by one of the interaction partners. Figure S8 shows that our results are robust to changes in the disparity threshold.

**Leadership** is a simple binary filter requiring that the participant who moved more in the first segment (*t*_1_ to *t*_2_) moved less in the second segment (*t*_2_ to *t*_3_). We refer to the participant who moved the most in the first segment max_*a*Pp*i, j*q_ Δ*x*_*a*_ *t*_1_, *t*_2_ as the *leader* and the participant who moved the most in the second segment max_*b*Pp*i, j*q_ Δ*x*_*a*_ (*t*_2_, *t*_3_)as the *follower*. Thus, successful pulls are defined as *a = b*, where the leader and follower are separate participants.

**Duration** was the final filter, where we required pulls to be at least 3 seconds in duration (since it takes 2.25 seconds to destroy a block). After all filters were applied, the average pull duration was 13.1 seconds ± 0.09 (SEM).

### Computational modeling

To better understand individual foraging decisions at a mechanistic level, we developed a computational modeling framework that sequentially predicts each block participants destroy based on different combinations of asocial and social features. We modeled the choice probabilities for each block destruction using a linear combination of block features **f** and regression weights **w** that represent the influence of each feature for participants’ block choices (Eq. 1). This was modeled using a categorical likelihood function with *B*_*k*+1_ possible outcomes (i.e., the number of remaining blocks available for choice at time *k +*1), with a softmax link function. Different models incorporate different sets of features in **f**, while some dynamic models additionally adapt the weights of specific features as a function of the elapsed time (at time *k*) since the last individually acquired reward or the last socially observed reward (using visual field analysis), depending on the model (see main text).

For interpretability of weight estimates and to allow for identical prior distributions, we *z*-standardized all block features within each choice, with the exception of block visibility, which was coded as a binary indicator. We also omitted the first choice in each round, since most features need to be computed with respect to some previous block destruction. Thus, we only started modeling from the second choice in each round, conditioned on the first choice. Furthermore, while all asocial features were included as predictors for each choice, the social features could be undefined for some choices if the conditions were not met (e.g., no visible players, or no visible and successful players). In these situations, the feature values were effectively set to 0 for all blocks.

All model weights were estimated in a hierarchical Bayesian framework with random effects accounting for differences in the importance of (asocial and social) features among individuals and experimental groups. The models were fit using Stan as a Hamiltonian Monte Carlo engine for Bayesian inference^73^, implemented in R v.4.0.3 through `cmdstanr` version 0.3.0.9. We used within-chain parallelization with `reduce_sum` to reduce model run times through parallel evaluation of the likelihood.

To minimize the risk of overfitting the data, we used weakly informative priors for all parameters. We used weakly informative standard normal priors centered on 0 for all weight parameters, exponential priors for scale parameters (with rate parameter *λ =* 1) and LKJ priors (with shape parameter *η =* 4) for correlations matrices^74^. To optimize convergence, we implemented the noncentered version of random effects using a Cholesky decomposition of the correlation matrix^75^. Visual inspection of traceplots and rank histograms^76^ suggested good model convergence and no other pathological chain behaviors, with convergence confirmed by the Gelman-Rubin criterion^77^ 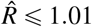 1.01. All inferences about weight parameters are based on several hundred effective samples from the posterior^78^. We provide additional details about some model features below.

### Block visibility

Since block visibility only captures a static representation of which blocks were visible at time *k*, we computed it with permissive assumptions. Specifically, we assumed no object or player occlusions (i.e., object permanence) and used only the horizontal component of their heading vector to avoid incorporating noise from vertical jitters. Visibility computations used the true horizontal viewing angle of 108.5 degrees, corresponding to the 16:9 aspect ratio monitors used in the experiment.

### Gaussian process for binary reward prediction

Gaussian processes^67^ provide a Bayesian function learning framework, which we use as a psychological model of reward generalization for predicting search behavior^37^. Gaussian processes are typically used to learn a function *f* : *𝒳 ⟶* ℝ^*n*^ that maps the input space *𝒳* (i.e., the field of destructible blocks) to real-valued scalar outputs, such as continuous reward values.

Here, we modify the Gaussian process framework to the binary classification case, where we want to make probabilistic predictions about whether destroying some block **x** will yield a reward *p*(*r =* 1|**x**). This can be described as a logistic sigmoid function *S*(·) of some real-valued latent variable *z*, such that 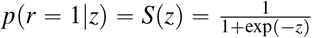. We set the prior mean 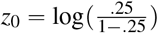 such that *p*(*r* = 1|*z*_0_) = 0.25, corresponding to the true prior probability of rewards. Thus, larger values of *z* correspond to higher-than-chance reward probabilities, while lower values correspond to lower-than-chance reward probabilities.

The latent variable *z* thus becomes the target of the Gaussian process posterior predictive distribution, computed for some location **x**_*_ P *𝒳* and conditioned on the past set of observations *𝒟*_*k*_ ={**X**_*k*_, **r**_*k*_}:

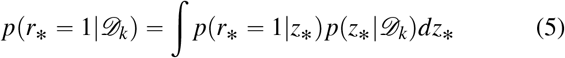

This exact integral in Eq. 5 is analytically intractable, but (i) assuming p(z_*_|*𝒟*_*k*_) is Gaussian distributed (using the Laplace approximation^67^; see below) and (ii) approximating *p*(*r*_*_ = 1|*z*_*)=_ (*S z*_*)_ with the inverse probit function^79,80^ Φ (*z*_*_), we obtain a tractable approximation.

We start by defining a posterior on the latent variable *z*_*_ corresponding to some unobserved block **x**_*_:

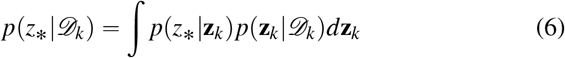

The first term *p* (*z*_*_| **z**_*k*_)is a Gaussian distribution that can be obtained using the standard GP posterior predictive distribution^67^, while *p*(**z**_*k*_ | *𝒟*_*k*_)is intractable. However, the Laplace approximation allows us to approximate the latter term using a Gaussian distribution:

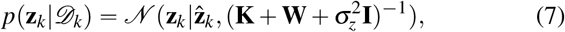

where 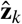 is the posterior mode, **W** is a diagonal matrix with diagonal elements 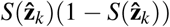 **K** is the *k × k* kernel matrix evaluated at each pair of observed inputs (see Eq. 12), 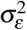 is the noise variance, and **I** is the identity matrix. We set 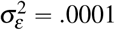 as in the environment generating process.The posterior mode 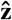 can be obtained iteratively:

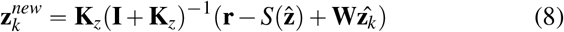

where 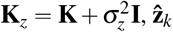 is the current estimate of the posterior mode, 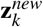 is the new estimate, and 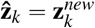 at convergence.

Eq. 6 can now be derived analytically as a Gaussian 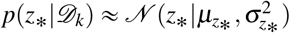, with mean and variance defined as:

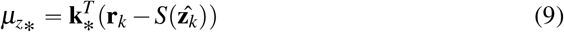

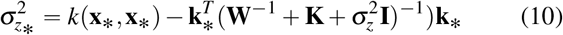

where **k**_*_ applies the kernel to the target **x**_*_ and all previously encountered observations **k**_*_=*k* (**x**_1_, **x**_*)_, …, *k* (**x**_*k*_, **x**_*)]_.

Lastly, we use the inverse probit function Φ (*z*_*_)as a common method^79,80^ for approximating the reward probability as a function of the mean and variance estimates described above:

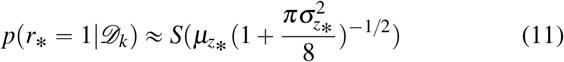

As a kernel function, we use the radial basis function kernel, which specifies that the correlation between inputs decays smoothly as a function of distance:

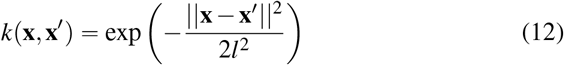

The degree of smoothness is controlled by the length scale *l*, which we set to 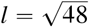. Note that this is equivalent to the *l =* 4 used to generate the environments, but accounts for the scaling of the coordinate system in the experiment, where each block has an empty tile on each side.

## Data Availability

All data^81^ collected from the experiment are available at github.com/charleywu/minecraftforaging.

## Code Availability

All code^81^ for running the experiment, analyzing the data, and the Unity simulations are available at github.com/charleywu/minecraftforaging.

## Acknowledgements

We thank Philip Jakob for technical support, Philipp Schwartenbeck for advice on cluster correction, and Ariana Strandburg-Peshkin and Damien Farine for publishing their code for analysing pull events. We also thank Ryutaro Uchiyama and Alexandra Witt for helpful feedback on a draft of the manuscript, and Deb Ain for copy editing. CMW is supported by the German Federal Ministry of Education and Research (BMBF): Tübingen AI Center, FKZ: 01IS18039A and funded by the Deutsche Forschungsgemeinschaft (DFG, German Research Foundation) under Germany’s Excellence Strategy–EXC2064/1–390727645. RHJMK and DD are funded by the DFG under Germany’s Excellence Strategy – EXC 2002/1 “Science of Intelligence” – project number 390523135. A pilot version of this experiment (with different data and design) was presented at the 43rd Annual Conference of the Cognitive Science Society^82^.

## Author contributions statement

CMW and RHJMK conceived the experiment, with feedback from DD, BK, and BM. CMW, DD, BK, and RHJMK performed the experiments. CMW, DD, and MHH analyzed the results. BK developed the visual field transcription method under the supervision of CMW and RHJMK. CMW developed the visualizations and wrote the first draft. All authors reviewed the manuscript.

