## Supplementary information for "Adaptive mechanisms of social and asocial learning in immersive collective foraging"

### Supplementary Videos

**Movie S1.** Tutorial. The original German text has been translated to English for better interpretability. `https://www.youtube.com/watch?v=QksKYOoElxg`

**Movie S2.** Bird's eye recreation of a group round with smooth rewards. `https://www.youtube.com/watch?v=vUHaAhjfFVo`

**Movie S3.** Screen capture of a group round with smooth rewards (corresponds to Supplementary Video 2). All in-game text was originally in German for all experiments, but have been translated here to English for interpretability. `https://www.youtube.com/watch?v=wyk7RbmHiok`

**Movie S4.** Screen capture of a group round with random rewards. `https://www.youtube.com/watch?v=mWe4CeLWdpg`

**Movie S5.** Automated transcription of visual field using Unity simulations. `https://www.youtube.com/watch?v=iSZ-ewpiZWI`

### Supplementary Methods

#### Rewards

Smooth environments resulted in higher absolute rewards in both solo (0.5 [0.4,0.5]) and group conditions (0.3 [0.3, 0.4]; Fig. S1a,c). Within smooth environments, participants performed better in solo than in group rounds (-0.1 [-0.2, -0.1]). However, this effect of solo vs. group condition disappears when we control for faster reward depletion in groups (Fig S1b) by computing normalized reward rate as `rewardRate/expectedRewardRate` (Fig. 2a), where `expectedRewardRate` is the marginal probability that any of the remaining blocks contains a reward. Thus, reward structure is the key driver of performance (Fig S1d).

#### Turning Angle

Using methods developed for studying foragers in naturalistic settings[1], we computed the average turning angle between block destruction events $\delta$. To do so, we first computed the *heading angle* $\theta$ based on the arctan of the displacement ratio between consecutive block destruction events $k$ and $k-1$:

$$\theta_k = \arctan\left(\frac{y_k - y_{k-1}}{x_k - x_{k-1}}\right). \quad (1)$$

We then compute the *turning angle* $\delta$ as the difference in heading angle, which is normalized to unit scale by dividing by $\pi$:

$$\delta_k = \frac{|\theta_k - \theta_{k-1}|}{\pi}. \quad (2)$$

In Figure S2b, we show how turning angle is influenced by whether or not the previous block yielded a reward. In general, turning angles were larger in smooth environments (0.04 [0.02, 0.06]; see Fig. S2c). The successful acquisition of reward resulted in larger angles in smooth (0.03 [0.02, 0.05]), but not random environments (-0.004 [-0.014, 0.007]). These results are consistent with the normative outcomes, since a greater degree of success-dependent adaptivity of turning angles corresponded to better performance in smooth but not random environments (i.e., more negative values in Fig. S4, corresponding to larger turning angles following success).

#### Social distance

We then computed the average pairwise distance between participants (Fig. 2c; Fig S5a). Solo rounds provide an asocial baseline by accounting for the influence of reward structure, which we calculated by simulating as if participants were on the same field.

### Foraging rate

We additionally looked at participant foraging rates, defined as the number of destroyed blocks per second. This analysis revealed greater selectivity in smooth environments, corresponding to a slower rate of blocks destroyed (-0.04, [-0.07, -0.02]; Fig. S5c-d). The selectivity of smooth environments was further amplified in group rounds (-0.03, [-0.06, -0.005]), where participants did not only need to contend with the structure of the environment, but also the structure and dynamics of social interactions.

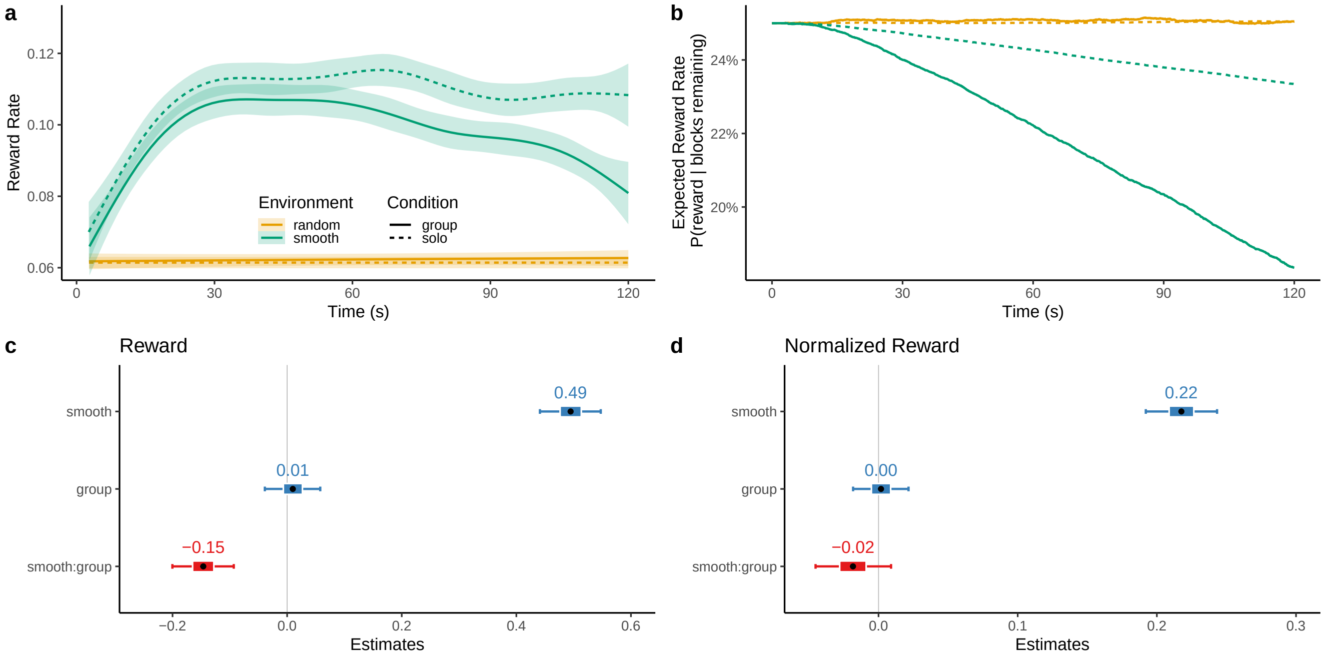


**Supplementary Figure S1. Reward**. **a**) Smoothed curves showing the average rate of rewards over time as a Generalized Additive Model (GAM). Ribbons indicate 95% CI. **b**) Expected reward rate over time (used to compute normalized reward rate; Fig. 2a), showing the probability that a randomly sampled block (from those remaining) contains a reward. Each line shows the aggregated mean, which only diminished in smooth environments (due to predictable rewards) and much faster in group rounds (due to more participants foraging for the same finite number of rewards). **c**) Coefficient plot of a hierarchical Bayesian Poisson regression showing (absolute) rate of rewards. Each dot is the posterior mean and error bars show the 95% CIs. **d**) When running a regression on the normalized rewards, only the effect of smooth environments remains.

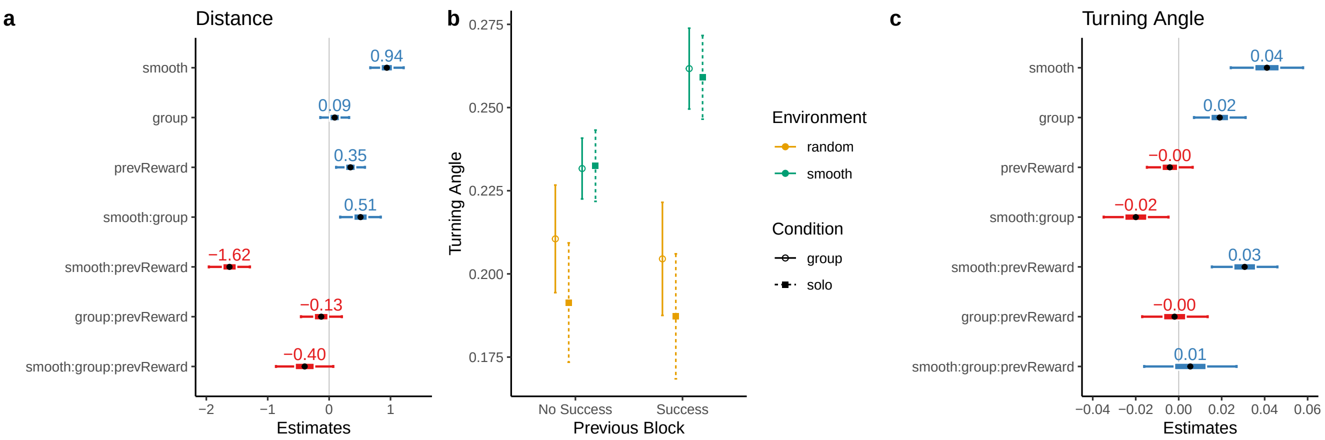


**Supplementary Figure S2. Adaptive foraging**. **a**) Coefficient plot of a hierarchical Bayesian regression on foraging distance (between blocks). Each dot is the posterior mean and error bars show the 95% CIs. **b**) Turning angle (Eq. 2) as a function of success. Each dot is a group mean with error bars indicating the SEM. **c**) Coefficient plot of a hierarchical Bayesian regression on turning angle.

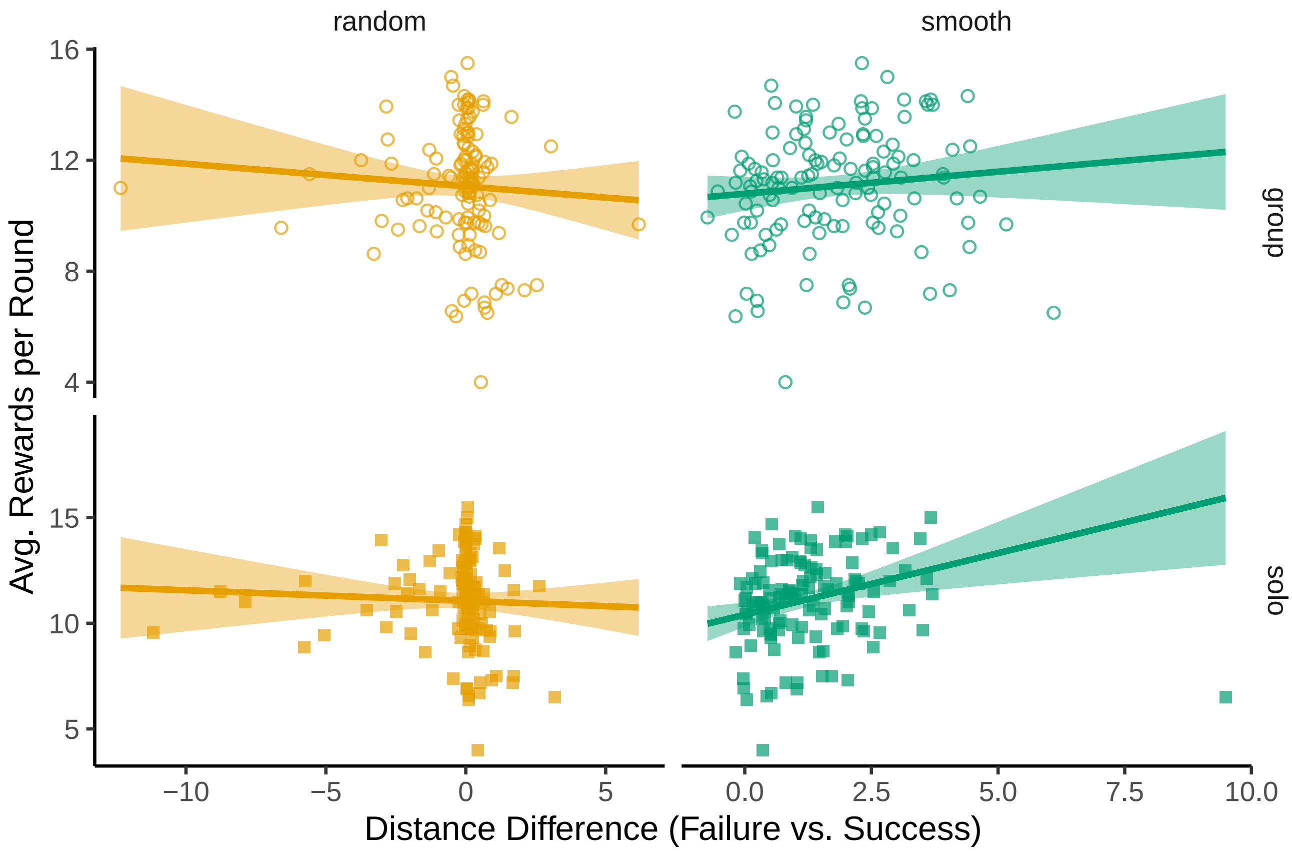


**Supplementary Figure S3. Adaptive foraging distance and performance**. Adaptive foraging distance is the mean difference of foraging distance (i.e., between consecutive blocks) after a failing to acquire a reward and after successfully acquiring a reward (larger values indicate a decrease of foraging distance after success). Rewards are averaged across the four rounds of each condition. Rewards were correlated to the adaptive distance in smooth (solo: $r_\tau = .20$, $p < .001$, $BF = 32$; group: $r_\tau = .18$, $p = .003$, $BF = 9.9$; Kendall's tau) but not random environments (solo: $r_\tau = -.00$, $p = .993$, $BF = .12$; group: $r_\tau = -.06$, $p = .328$, $BF = .19$). Each dot is a participant, while the lines and ribbons are a linear regression (± 95% CI). A single outlier in smooth:solo was omitted from the linear regression line (around 10 on the x-axis), but not from the rank correlations.

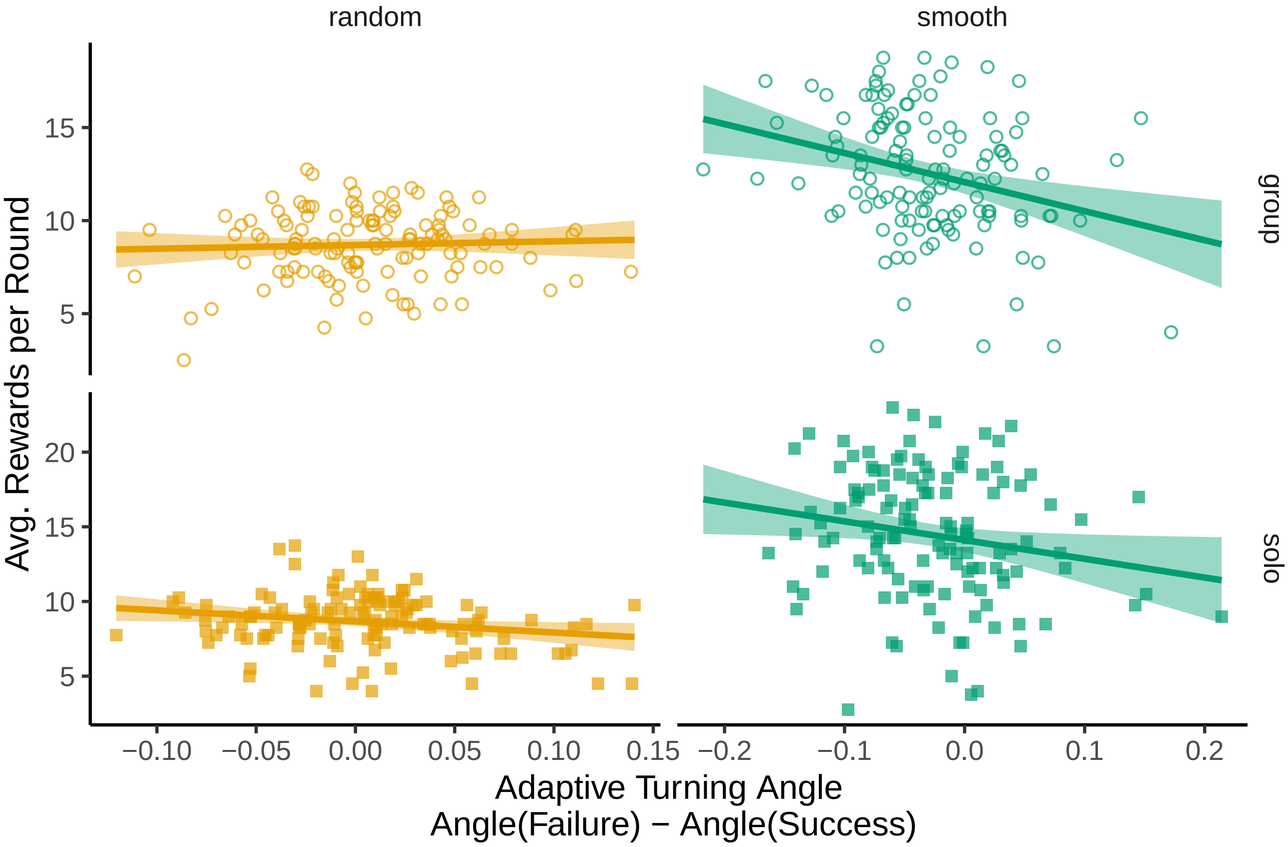


**Supplementary Figure S4. Adaptive turning angle and performance**. Turning angle (Eq.2) is commonly used to measure foraging trajectories in the context of area restricted search[1,2], where greater angles correspond to more local search. We measured the mean difference of turning angles after a failing to acquire a reward vs. after successfully acquiring a reward (more negative values indicate a larger increase of turning angle after success, i.e., more local search following success). Rewards are averaged across the four rounds of each condition. Rewards were correlated to the adaptive turning angle in smooth (solo: $r_\tau = -.14$, $p = .020$, $BF = 1.8$; group: $r_\tau = -.18$, $p = .003$, $BF = 10$) but not random environments (solo: $r_\tau = -.08$, $p = .176$, $BF = .30$; group: $r_\tau = .01$, $p = .892$, $BF = .12$). Each dot is a participant, while the lines and ribbons are a linear regression ($\pm$ 95% CI).

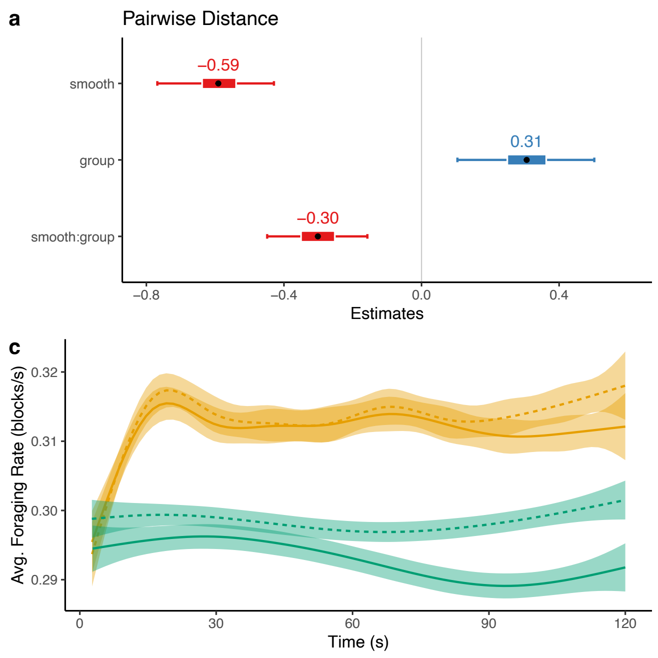


**Supplementary Figure S5. Pairwise distance and foraging rates**. **a**) Social distance regression coefficients. Participants were closer together in smooth environments and solo conditions. In random environments, participants in groups avoided each other compared to the solo condition. **b**) Visibility regression coefficients. Participants observed each other more in smooth environments and marginally less in the group condition for random, but not smooth environments. **c**) Foraging rate (i.e., the number of blocks destroyed per second) plotted as smooth conditional means. **d**) Regression coefficients. Participants had a lower foraging rate in smooth than in random environments, which was further amplified in group rounds.

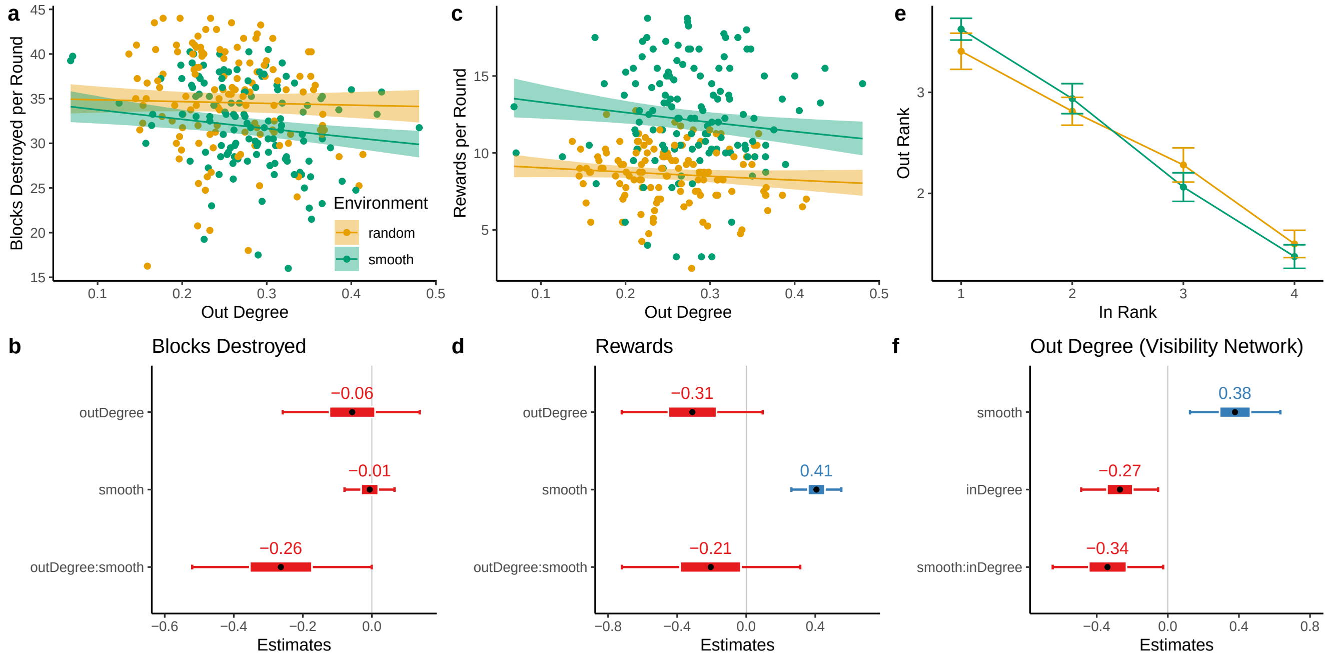


**Supplementary Figure S6. Relationships between environment, out-degree, in-degree, blocks destroyed and reward rate in groups**. **a-b**) In smooth environments, participants with a higher out-degree (i.e., observing other players) destroyed fewer blocks (Poisson regression coefficients shown in panel **b**). **c-d**) This did not translate into an effect on the reward rate. **e**) Rank ordering participants in each group according to their in- and out-degree showed a negative correlation between participants' in- and out-degree. **f**) Regression coefficients predicting out-degree. Smooth environments increased out-degree, while higher in-degree decreased out-degree, with a reliably stronger negative effect in smooth environments.

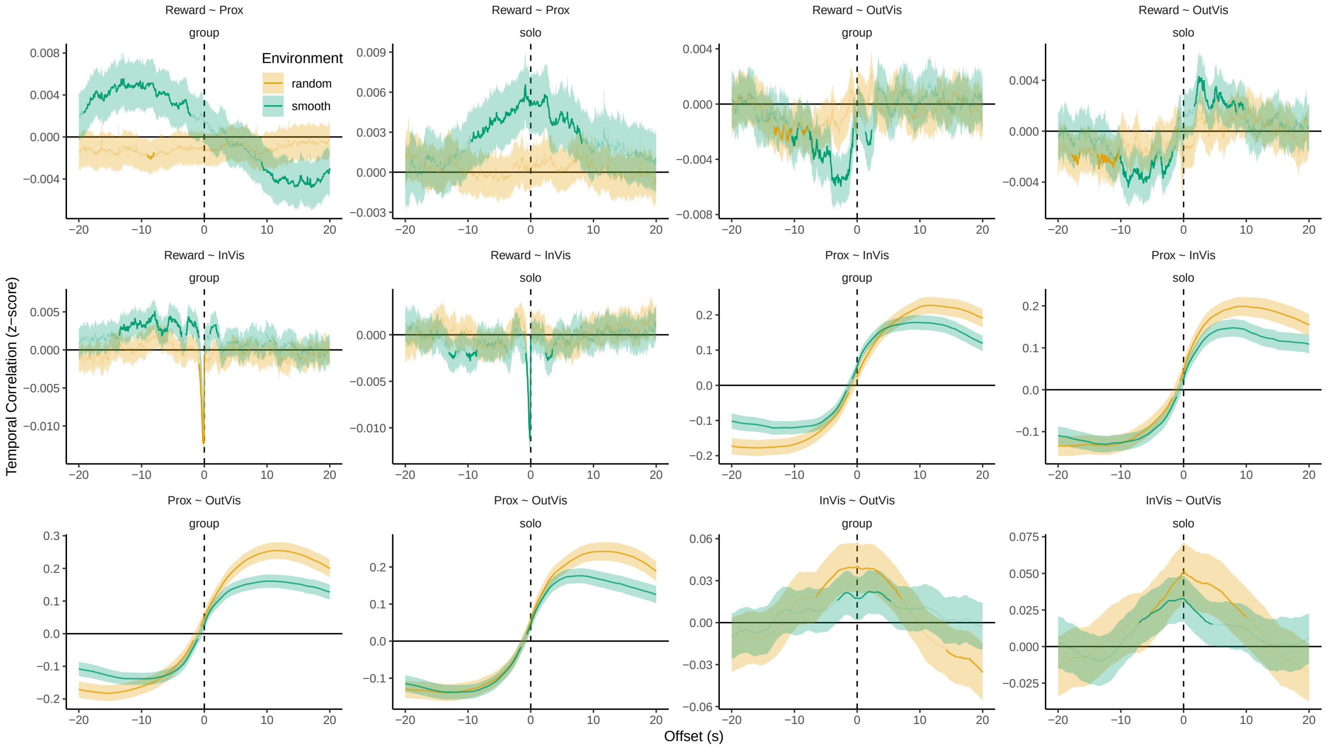


**Supplementary Figure S7. Temporal dynamics**. Full set of temporal dynamic analyses, including solo rounds. Lines show the group means and ribbons show the 95% CI. Bold lines indicate significant clusters that survived the permutation analysis (see Methods).

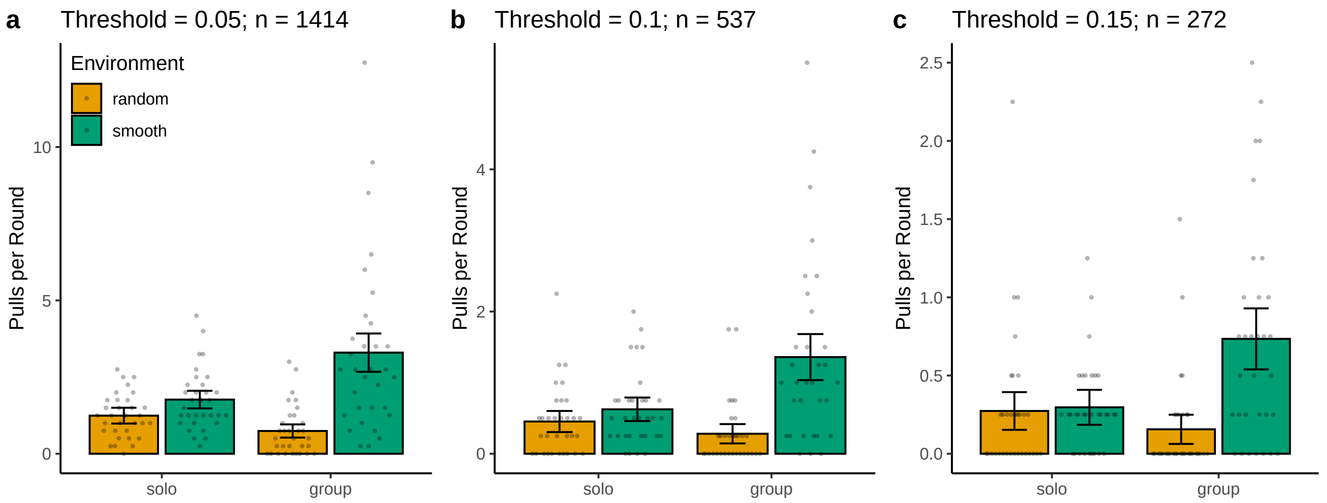


**Supplementary Figure S8. Sensitivity analysis of the pull analysis. a-c**) Independent of the exact disparity threshold used (0.05, 0.1, 0.15; see Methods), the number of pull events decreased from solo to group rounds in random environments, and increased from solo to group rounds in smooth environments. Dots show individual sessions, while bars show the aggregate mean and error bars are the 95% CI.

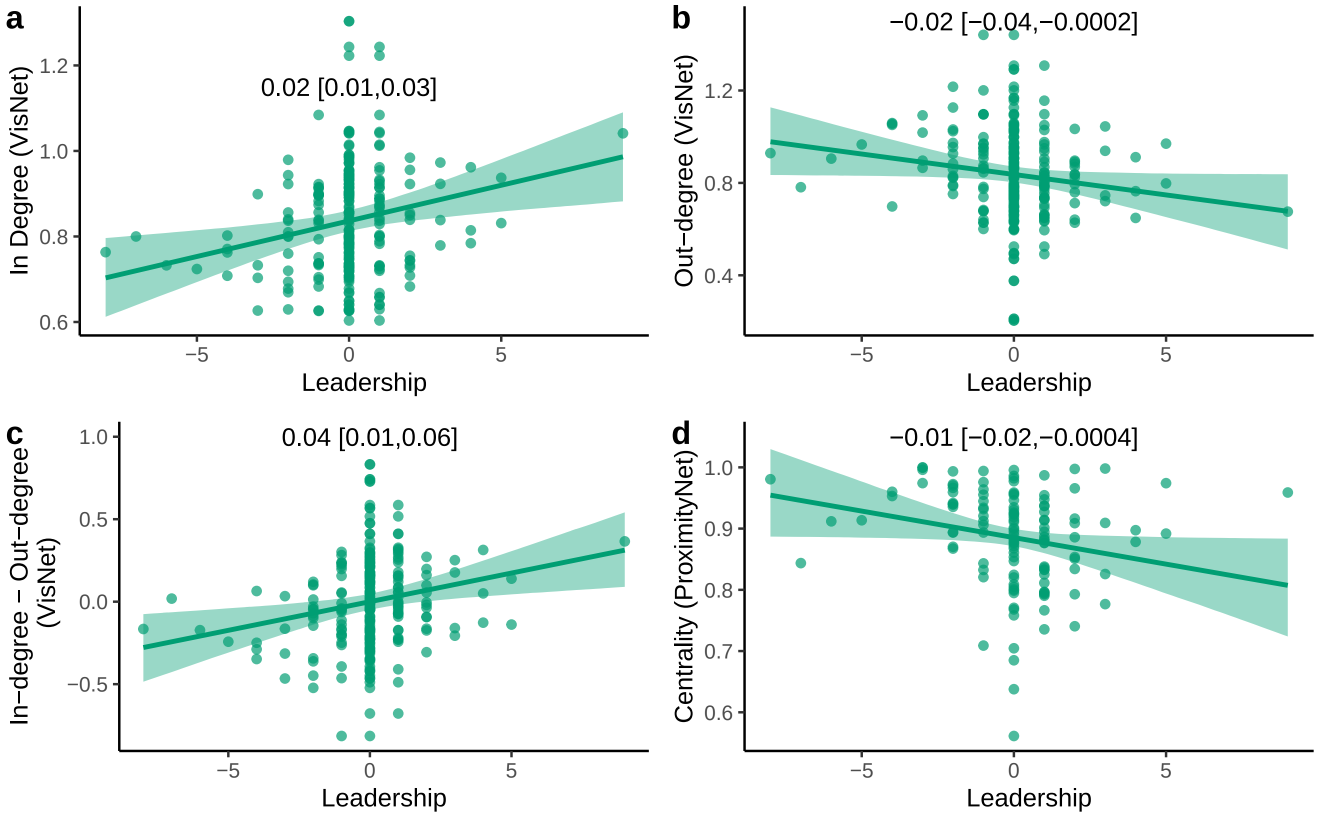


**Supplementary Figure S9. Relating leadership to visibility (VisNet) and proximity networks (ProximityNet)**. Each dot is a participant, with the line (and ribbon) showing the mean (±95% CI) of a mixed-effects regression, with the fixed effect reported above. For interpretability, we compute leadership only from group rounds in smooth environments. **a**) Participants with a higher leadership score were observed more (i.e., higher in-degree), and **b**) observed others less (i.e., lower out-degree). **c**) Leadership score also predicted the difference between in-/out-degree, and **d**) high leadership score also predicted lower spatial centrality, suggesting leaders were at the frontiers of the group.

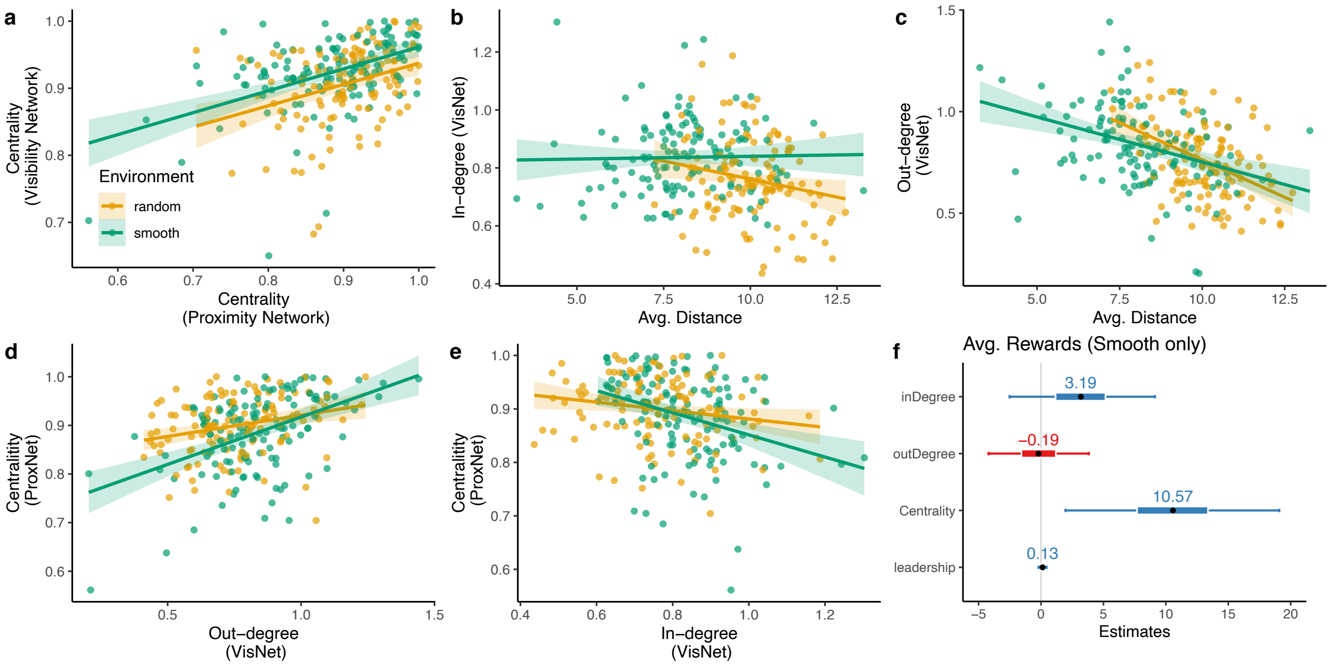


**Supplementary Figure S10. Consistency of visibility and proximity networks and their relationship to performance (in group rounds).** Each dot is a participant, with the line and ribbon showing a linear regression. **a**) Eigenvector centrality was consistent across visibility and proximity networks. **b**) A player's in-degree was unrelated to their average spatial distance to other players in smooth environments, and negatively correlated in random environments. **c**) Average distance to other players was always negatively correlated to out-degree. **d**) Out-degree was positively correlated with centrality, with a stronger effect in smooth environments. **e**) In-degree was negatively correlated with centrality in both environments. **f**) Bayesian mixed-effects regression predicting the influence of social network statistics on reward (smooth rounds only).

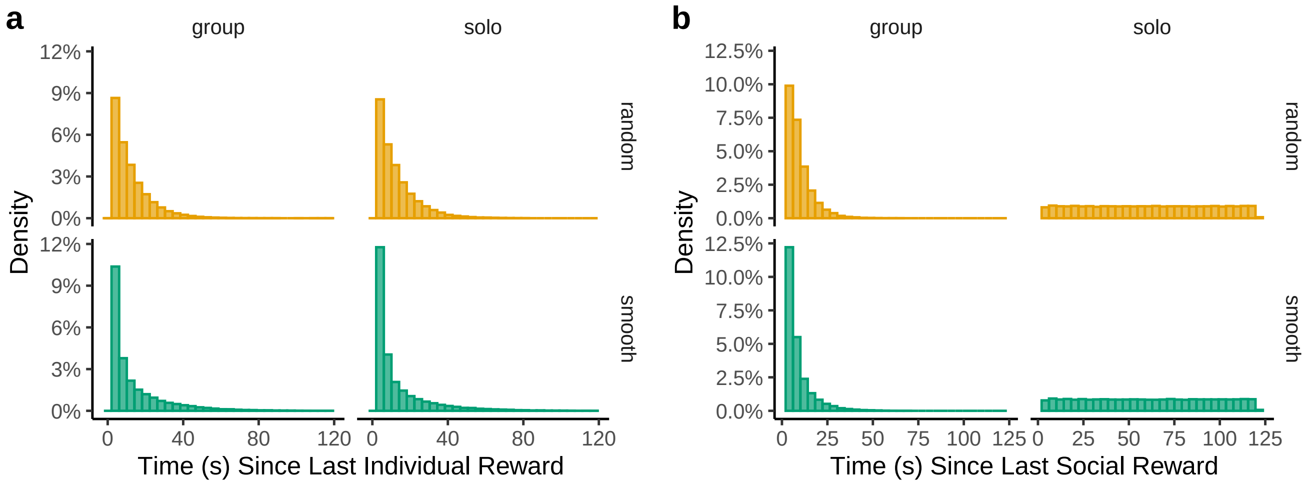


**Supplementary Figure S11. Histograms of elapsed time between previous individual/social reward events**, computed at each block destruction. These values are used in the adaptive models. Note that solo rounds do not contain social reward information.

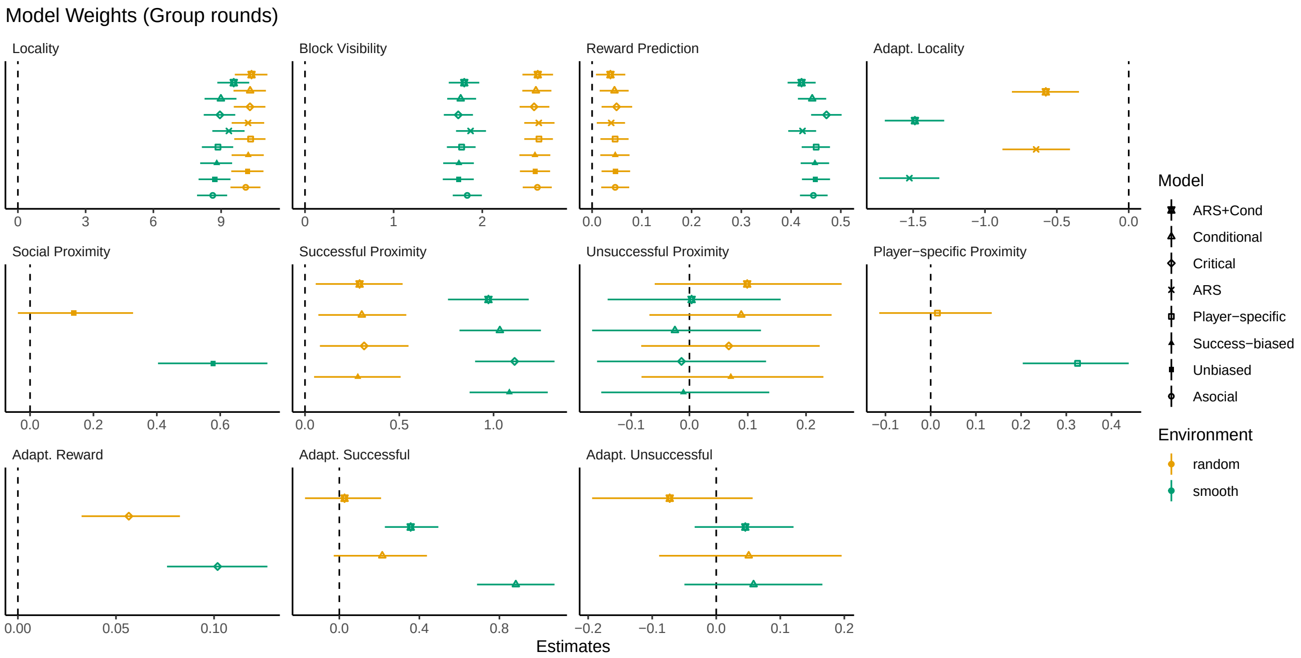


**Supplementary Figure S12. Weights for all models in group rounds**. Symbols show the posterior mean and error bars the 95% HPDI.

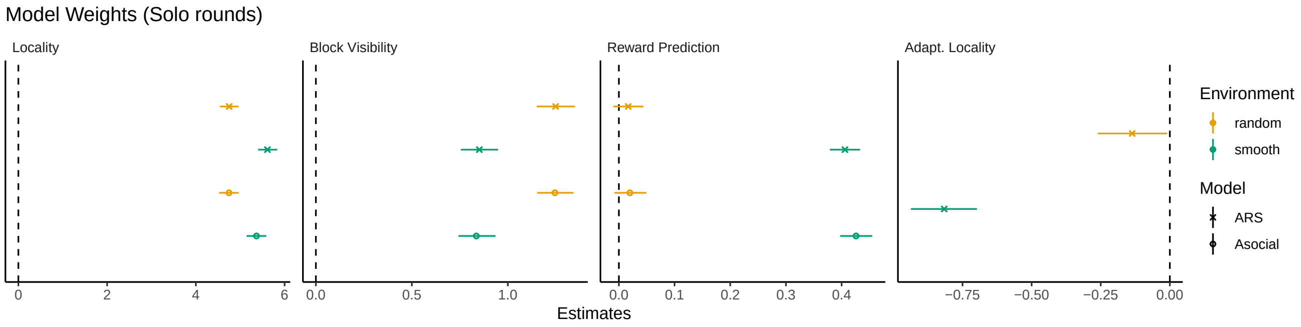


**Supplementary Figure S13. Weights for all models in solo rounds**. Symbols show the posterior mean and error bars the 95% HPDI.

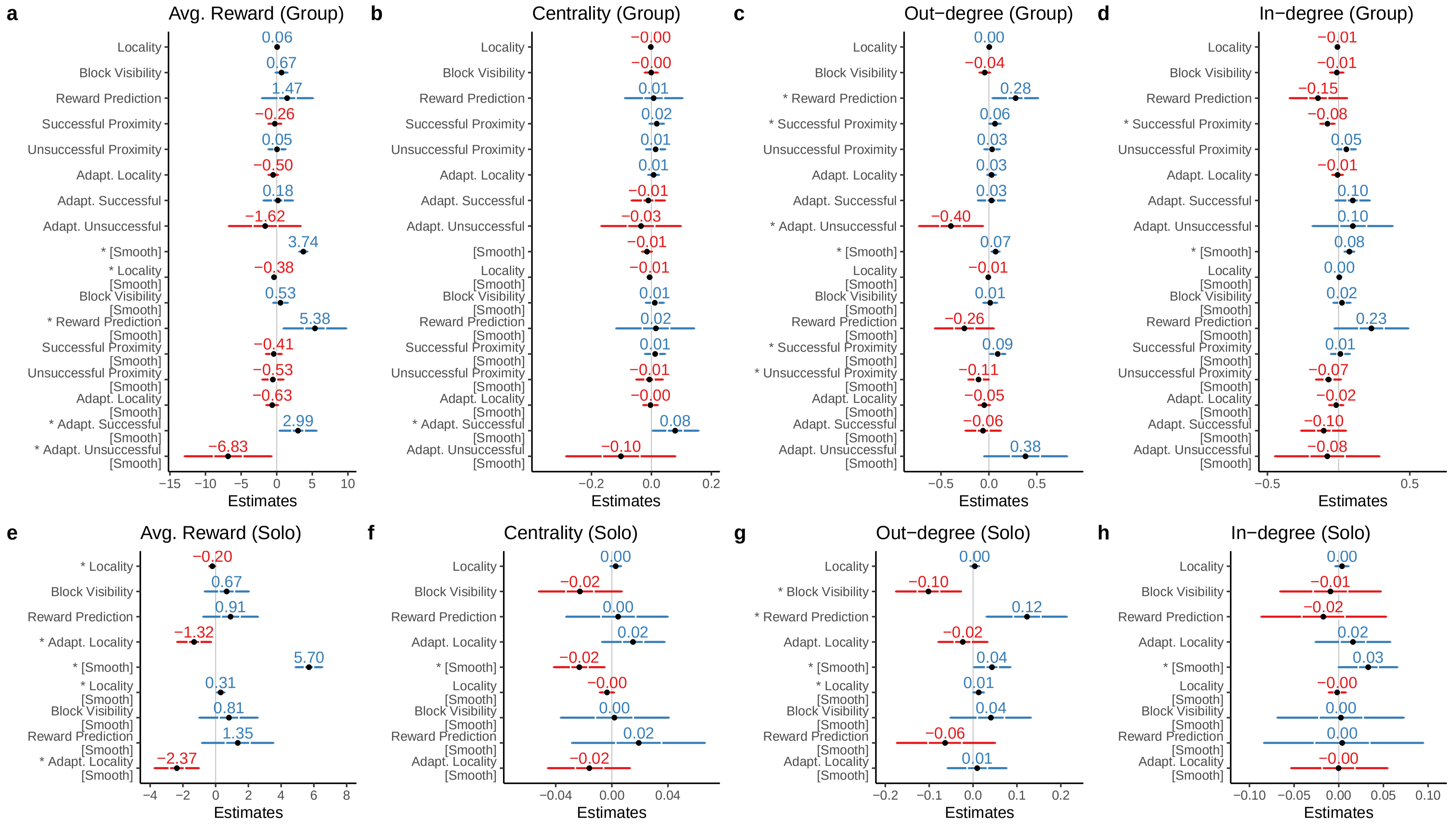


**Supplementary Figure S14. Regression coefficients of model weights on behavioral variables**. **a-d** are results using the winning ARS+Cond model using data on group rounds, while **e-h** are results for the winning ARS model on solo rounds. All models are hierarchical Bayesian regressions, predicting either average reward (**a**,**e**), centrality (proximity network; **b**,**f**), out-degree (visibility network; **c**,**g**), or in-degree (**d**,**h**), using individual estimates of model weights and their interaction with environment type (random by default, and smooth when indicated). ∗s on the y-axis labels indicate coefficients where the 95% HPD does not overlap with 0.

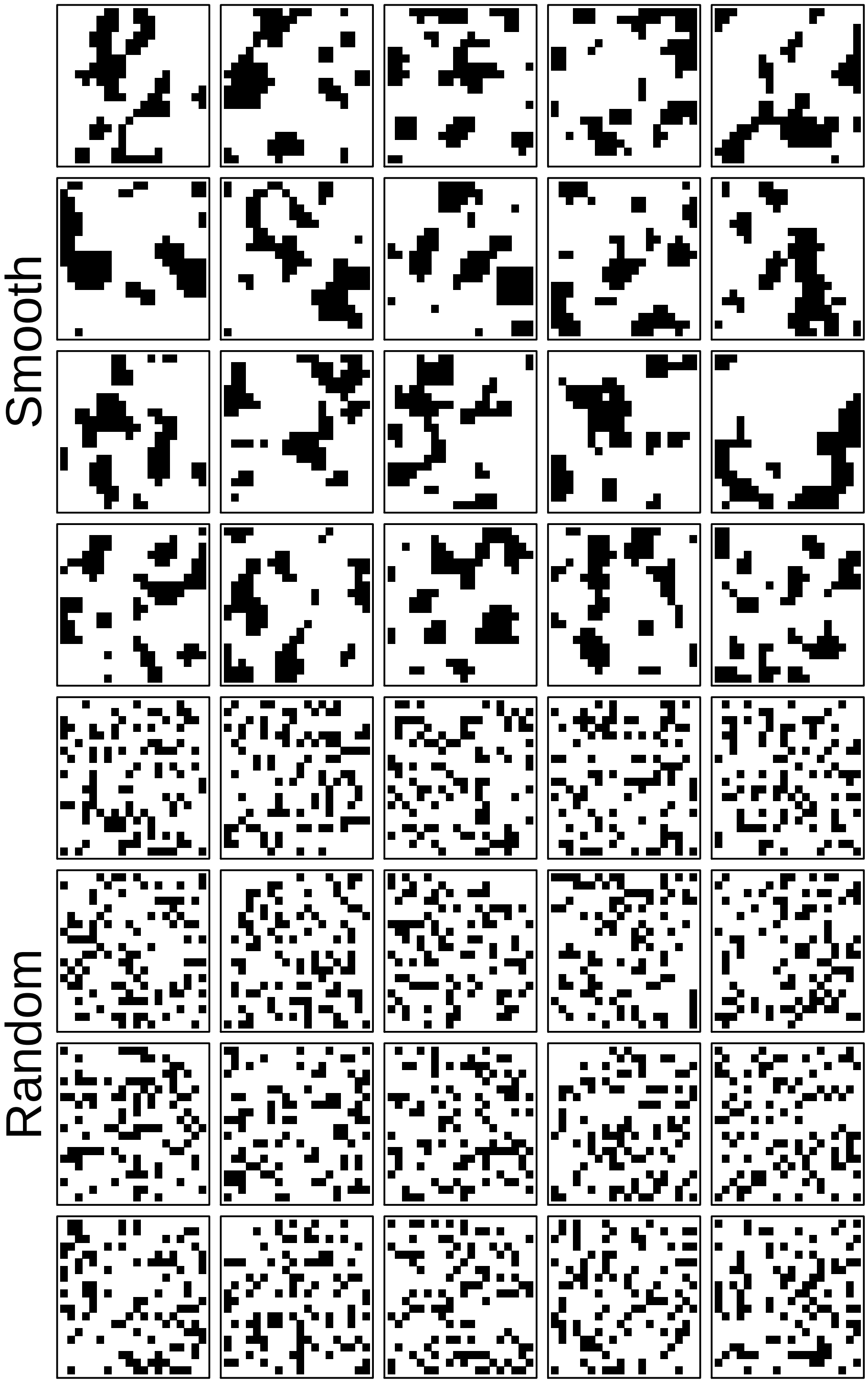


**Supplementary Figure S15. Reward distributions used in the experiment**. Black squares represent blocks containing a reward, while white squares represent boxes containing no reward. Note that these plots omit the spacing between resource blocks in the experiment for readability.
